# The folding, dimerization and allosteric landscapes of the ALS protein SOD1: a comprehensive mutational atlas

**DOI:** 10.64898/2026.07.15.738613

**Authors:** Tomás Quiroga, Laz Ashcroft, Lauren Rice, Defne Boratav, Mariano Martín, Juan Francisco Vázquez-Costa, Bryony A Thompson, Luke McAlary, Benedetta Bolognesi

## Abstract

Mutations in superoxide dismutase 1 (SOD1) can disturb monomer folding and dimerization, ultimately leading to the development of amyotrophic lateral sclerosis (ALS). Genotype-phenotype knowledge of SOD1 monomer folding and dimerization is largely incomplete, impeding our ability to treat and understand SOD1-ALS. To address this issue, we performed multidimensional deep mutational scanning of ∼6,000 SOD1 variants (amino acid substitutions, insertions, deletions), quantifying folded monomer abundance, and measuring wild-type-variant heterodimerization. Importantly, both abundance and heterodimerization capture pathogenicity, with heterodimerization being a slightly better classifier of pathogenic variants. Notably, ∼80% of pathogenic variants and ∼70% of variants of uncertain significance in SOD1 perturb both monomer abundance and heterodimerization, pointing to protein destabilization as a major driver of SOD1-ALS. Orthogonal validation in mammalian cells shows that protein abundance strongly correlates with other disease-relevant phenotypes, including protein aggregation and toxicity. Although monomer abundance and heterodimerization are strongly coupled, residual effects identify variants that perturb the heterodimer beyond their impact on monomer folding. Mapping residual effects uncovers potential allosteric sites that modulate dimer stability independently of monomer folding, revealing mechanistic insights and potential therapeutic entry points to stabilize SOD1 as a dimer. Together, these results demonstrate that multidimensional phenotype variant mapping improves mechanistic understanding of SOD1 variants and clinical variant interpretation, while uncovering structural targets for SOD1-ALS treatment.

## INTRODUCTION

Superoxide dismutase 1 (SOD1) is a highly stable, abundant, and ubiquitously expressed protein that protects cells from oxide radicals. To date, hundreds of SOD1 variants have been identified in the population that can be causative for the devastating neurodegenerative disease amyotrophic lateral sclerosis (ALS)^1,2^. Initially, SOD1-ALS variants were hypothesized to cause disease by ablating SOD1 enzymatic function^3^, however, evidence now shows that a gain-of-toxic function through misfolding is the main driver of SOD1-ALS^2,4^. Importantly, this pathogenic mechanism is characteristic of both SOD1-associated familial (fALS) and sporadic (sALS) forms of disease, where SOD1 variants are present in ∼20% and in ∼2% of cases, respectively^5,6^.

Structurally, SOD1 is composed of 153 amino acids that fold into a Greek key β-barrel with eight β-strands and eight loops that collaborate to bind zinc and copper, and form an intramolecular disulfide bond^2,7,8^. Loop IV is the zinc-binding loop and is responsible for coordinating zinc to stabilize the protein, meanwhile loop VII is the electrostatic loop and is responsible for guiding oxide radicals to the enzymatic cleft^2,9^. Maturation of nascent SOD1 by binding zinc, copper, and forming the disulfide is followed by homodimerization, resulting in an extremely stable protein^10,11^.

Whilst wild-type (WT) SOD1 is highly stable, at least some SOD1-ALS variants have been shown to differentially reduce folding stability in a variant-specific manner^12–15^, ultimately leading to misfolding and subsequent protein aggregation^14,15^. These variants can occur in multiple forms including the most common amino acid substitutions, as well as premature stops that lead to truncation^16^ and inframe amino acid deletions^17^. The effect of SOD1 variants on SOD1 structure is often multifactorial, with most variants affecting several features at once^12,18^. For example, the A4V (p.Ala5Val) variant can impede monomer folding and can also disrupt dimerization of mature monomers^12,19,20^. In contrast, the H46R (p.His47Arg) variant prevents copper binding^21^ but only minimally affects stability of either the monomer^22^ or dimer^23^. In line with this, the phenotypes observed in SOD1-ALS patients are also variant dependent^24–26^. Patients with the A4V variant display rapid disease progression (<1 year survival post-diagnosis)^1,27^, whereas patients with the H46R variant display a substantially slower progression (∼17 years survival post-diagnosis)^28^. In addition, heterodimerization of variants with WT SOD1 has been suggested as a contributor to SOD1 ALS as the stability of five variant–WT SOD1 heterodimers was shown to strongly correlate with patient survival time^29,30^. Computational methods have been employed to assay multiple characteristics of hundreds of SOD1 variants but lack strong experimental supporting data^31,32^. Additionally, *in vitro* and *in vivo* examination has been more accurate at measuring variant effects but has focused on a limited number of variants due to throughput^33,34^. Attempts to map the genotype-phenotype relationship of SOD1 variants have focused on activity and protein levels, the former of which is not associated with disease^35^. Multiplexed abundance assays, where abundance is a proxy measure of protein stability, have provided a comprehensive map of the effects of missense variants on SOD1 protein levels, establishing protein destabilization as an important contributor to pathogenicity^35^. More recently, small-scale biophysical studies have also examined the effects of mutations on protein stability at individual residues, such as G93^36^. However, because SOD1 variants can perturb multiple aspects of protein behavior, it remains unclear how abundance relates to other disease-relevant phenotypes such as dimerization, aggregation, and cell survival, and a comprehensive mapping integrating these features has yet to be performed.

Taking the above into account, the genotype-phenotype relationship for SOD1-ALS remains mostly unresolved. Indeed, increasing use of next-generation sequencing in the clinic to identify new *SOD1* variants has revealed substantial gaps in our understanding of how sequence variation translates into downstream phenotypes^5^. Currently, of all the more than 180 mutations reported for SOD1, ∼40% are missense variants of uncertain significance (VUS) or with conflicting classifications of pathogenicity, and ∼60% are well-classified as pathogenic^37,38^. However, quantitative evidence linking molecular mechanisms to disease exists only for a few variants^5^. This issue prevents our ability to better understand disease mechanisms, design clinical trials, and identify at-risk patients for early intervention.

Considering this, we set out to directly address the effect that SOD1 variants have on folding stability and heterodimer formation. Cell-based multiplexed assays of variant effects (MAVEs)^39,40^ exploit cell machinery to report mutational impact on specific phenotypes and to generate comprehensive atlases of variant effects^41^. In this work, we experimentally assayed ∼6,000 SOD1 variants, including single amino acid substitutions, insertions, and deletions, in two deep mutational scanning (DMS)^42^ assays. In one assay we measure folded monomer abundance and in the other we quantify heterodimerization of the mutant SOD1 with WT SOD1^43^. These assays allowed us to create complete variant effect maps for SOD1 folding and heterodimerization. We further used this quantitative information to clinically classify SOD1 variants of uncertain significance (VUS) more accurately than previous work^35^, to identify variants that cause ALS independently of protein destabilization and to reveal potential allosteric sites that are distal from but contribute to dimer formation. Overall, this data contributes to a greater understanding of SOD1-ALS mechanisms and supports the usage of multidimensional DMS in improving variant classification.

## RESULTS

### Substitutions, insertions, and deletions have structure-dependent effects on SOD1 abundance

To quantify the effects of mutations on protein abundance, we synthesized a mutational library of ∼6,000 SOD1 variants which were all assayed through phenotypic selection and measured with deep sequencing (Fig. 1A, see Methods). We employed an abundance protein fragment complementation assay (abundance PCA)^43^ that uses dihydrofolate reductase (DHFR) complementation in the presence of methotrexate (MTX). Abundance PCA (aPCA) is used as a proxy of SOD1 foldedness, where the levels of the SOD1-DHFR fusion change due to unfolded or misfolded proteins being degraded, resulting in variants that lower SOD1 stability leading to impaired growth and a lower abundance (Fig. 1B). Abundance PCA replicates are highly correlated (median of Pearson’s R=0.96) (Fig. 1C and Suppl. Fig. 1C) and we obtained confident abundance scores for 5929 variants including single amino acid substitutions (n=2889), single amino acid insertions (n=2758), single amino acid deletions (n=153), synonymous (n=87) and premature stop (n=42) variants (Fig. 1D). The dynamic range of the assay clearly discriminates scores of premature stop and synonymous mutations, while scores of both missense and indel mutations are spread all along the dynamic range (Fig. 1D). Overall, we find a proportion of substitutions behave as WT-like (∼50%), while for insertions and deletions this percentage is lower (30% and 25%, respectively). The percentage of low abundance variants is smaller for substitutions (31.3%) but larger for insertions and deletions (46% and 38.5%, respectively), suggesting that indels have a more detrimental effect on protein abundance than substitutions, and agrees with previous work showing the dominant impairment effect of indels on protein stability^44^. Across the three types of mutations, deletions contain the highest proportion of stop-like mutations (36%), followed by insertions (22%) and substitutions (13%) (Fig. 1E).

**Figure 1.**
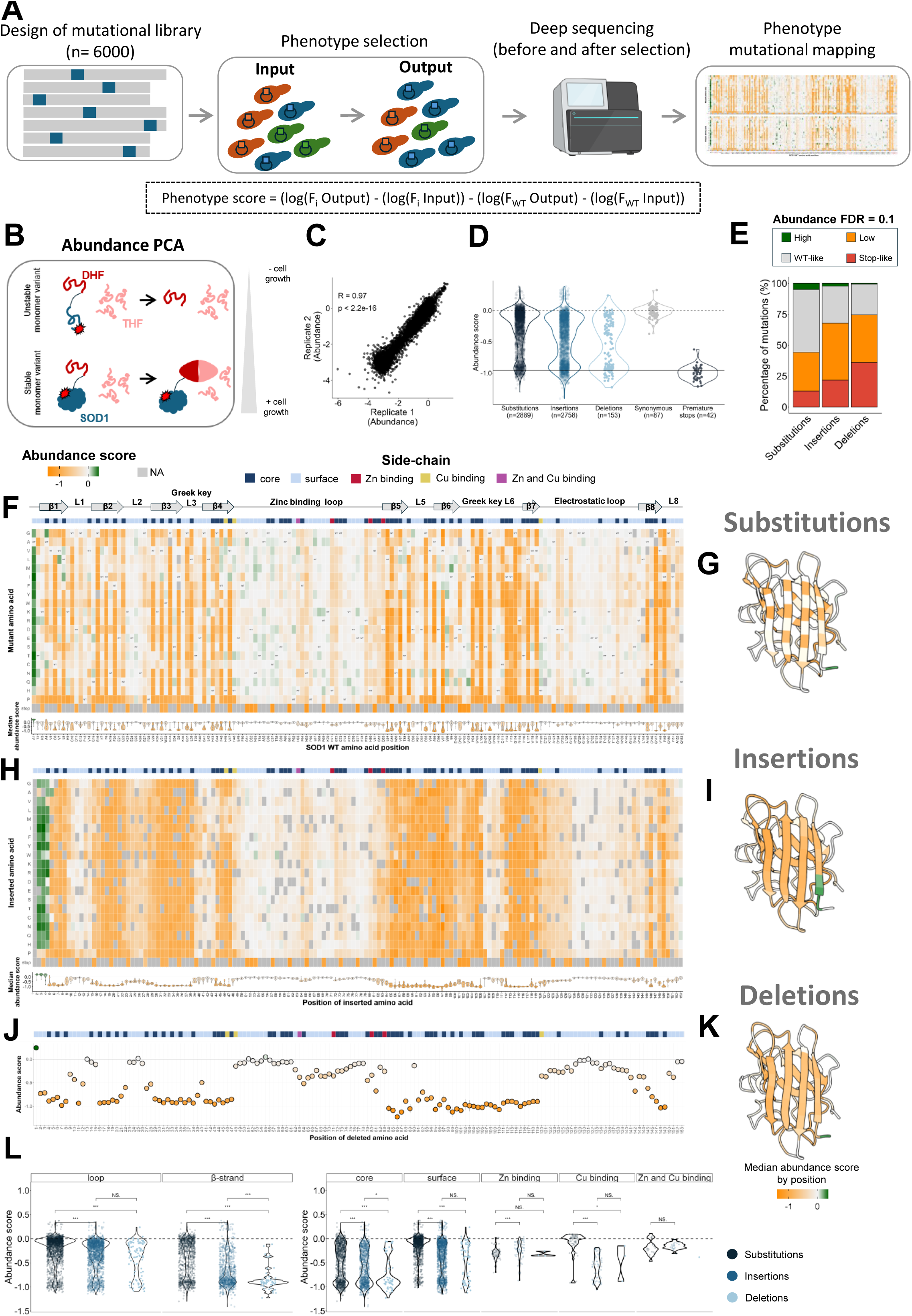
AbundancePCA of SOD1 variants reveals the basis of structural stability. **(A)** Deep mutational scanning workflow. A library of ∼6,000 SOD1 variants is expressed in yeast, and the DNA is extracted before (input) and after (output) selective conditions. The extracted DNA in both conditions is sequenced and variant scores are used for variant effects analysis **(B)** AbundancePCA (aPCA) workflow. Each SOD1 variant is fused to a fragment of DHFR (DHF), while the remaining DHFR fragment (THF) is over-expressed. **(C)** Scatter plot of aPCA replicates 1 and 2. Pearson’s correlation coefficient and p-value are indicated **(D)** Normalized score distribution of each variant type of the library for aPCA. The dashed line represents the WT-like score and the solid line represents the stop mutations mode. **(E)** Percentage of variants of each abundance FDR category for amino acid substitutions, insertions, and deletions. Abundance mutational maps for **(F)** amino acid substitutions, **(H)** insertions, and **(J)** deletions. Orange: decreased abundance; grey: WT-like abundance; green: increased abundance. The bar over each map represents the side-chain orientation of each position (darkblue: core residues, rSASA dimer < 25; lightblue: surface-exposed residues (rSASA dimer > 25); red: zinc-binding residues; purple: copper-binding residues). SOD1 3D monomer structures (PDB: 2V0A) colored by the median abundance scores per positions of substitutions **(G**), insertions **(I**), and deletions **(K).** (**L)** Violin plots showing the abundance scores distribution of mutations in loops, β-strands, and according to the side-chain. Significance is calculated with the Mann-Whitney U test: p>0.05 (NS, non-significant), p<0.05 (*), p<0.01(**), p<0.001(***).

SOD1 forms an 8-stranded Greek-key β-barrel that is stabilized by internal hydrophobic contacts, which is likely susceptible to variant-induced destabilization. Meanwhile, the long zinc-binding (Loop IV) and electrostatic (Loop VII) loops are less likely to result in destabilization when mutated and require zinc and copper co-factors to fold. In line with this, mapping substitutions to the SOD1 structure reveals that 58% of variants across the eight β-strands have a detrimental impact by lowering protein abundance and thus they likely disrupt SOD1 structural integrity, whereas loop variants are far less likely to lower protein abundance (Fig. 1F and G). This is mirrored for both insertions (Fig. 1H and I) and deletions (Fig. 1J and K), strongly suggesting that the SOD1 β-barrel core is highly sensitive to variant-induced destabilization. Supporting this, there is a bimodal distribution in the abundance scores of substitutions (Fig. 1D) which corresponds with side-chain orientation; namely, substitutions in surface-exposed residues (relative solvent-accessible surface area; rSASA > 25, Methods)^45^ result in WT-like scores whilst buried residues (rSASA < 25) result in negative scores (Fig. 1L).

Loop variants have a range of effects where short loops (I, II, III, V, VI and VIII) are more sensitive to mutation than long loops (zinc-binding loop IV and Electrostatic loop VII) which are likely to allow some structural rearrangement upon side-chain and backbone alterations (Fig. 1F, H, J and L and Suppl. Fig. 2B). Loops III and VI of the Greek-key structure show markedly lower tolerance to substitutions and deletions, whereas loops II and V are more substantially affected by insertions (Fig. 1F, H, J and Suppl. Fig. 2B). Greek-key loops III and VI form the β-barrel turns that connect opposite sides of the barrel, and as such they may have a more relevant role in maintaining or forming the native SOD1 fold.

During SOD1 maturation, each monomer binds one atom of zinc that confers substantial structural stability to the nascent protein, and one atom of copper that contributes less to folding stability^2^. The residues involved in metal binding form the catalytic site of SOD1. In line with this, all mutations of zinc-binding residues in our dataset are deleterious (Fig. 1L). More specifically, mutations of the zinc-binding residues H71 and D83 show a moderate detrimental effect. In contrast, while deletion of H80 leads to moderate destabilization, some substitutions of H80 are well tolerated (Fig. 1F, 1J and Suppl. Fig. 2C). This suggests that H71 and D83 are more important to coordinating zinc than H80. Interestingly, adjacent residues to zinc-binding residues are also detrimentally affected (G72, R79, and G82) in substitutions and indels, suggesting a potential role of these residues when it comes to zinc binding. For the discrete copper binding residues H46 and H120, we observe similar effects on abundance. Both H46 and H120, one with an exposed side chain and the other located inside the Electrostatic loop, tolerate substitutions, but not insertions or deletions (Fig. 1F, H, J and L). Finally, H63, a residue that coordinates both zinc and copper, is moderately sensitive to substitutions, deletions, and insertions (Fig. 1F, H and J).

When considering amino acid type and class, we found that substitutions at glycines, hydrophobic, and aromatic residues have the most detrimental impact (Suppl. Fig. 2D) on SOD1 abundance. Similarly, insertions at and deletions of hydrophobic and aromatic residues result in the lowest protein abundance. The detrimental impact of deleting hydrophobic and aromatic residues suggests an important role of these residues in β-barrel core stabilization (Suppl. Fig. 2D). Meanwhile, in SOD1 residues corresponding to the β-barrel, substitutions to and insertions of proline have the most detrimental effect on SOD1 abundance (Suppl. Fig. 2E), which matches previous mammalian cell measurements of proline substitutions in β-strands of SOD1^46^. In addition, we also observe that substitutions and deletions at the N-terminus of SOD1 increase the abundance of SOD1 relative to WT (Fig. 1F and H). This stabilizing effect occurs until position 4 when inserting amino acids (Fig. 1H). However, substitutions to proline in the first position and insertions of proline in positions 2 to 4 lead to destabilization (Fig. 1F).

By comparing the impact of mutations to the same amino acid across mutation classes, we find a high correlation for substitutions versus insertions (Pearson’s R=0.74, p=2e-04), where substitutions to/insertions of proline and tryptophan have the most deleterious effect (Suppl. Fig. 2F). When comparing instead the impact of mutations from the same amino acid, the correlation is high between insertions and deletions (Pearson’s R=0.69, p=1.6e-03), with tryptophan, isoleucine and valine being equally affected when deleted or when inserting residues (Suppl. Fig. 2G), and moderate between substitutions and deletions (Pearson’s R=0.57, p=1.3e-02) and between substitutions and insertions (Pearson’s R=0.58, p=1.1e-02) (Suppl. Fig. 2H and I).

Importantly, the correlation scores of insertions and deletions is high (R=0.69, p=8.6e-23), suggesting that regardless of the backbone perturbation (i.e., inserting or deleting one amino acid) the mutational impact will be similar (Suppl. Fig. 2J).

### SOD1 abundance measured in yeast is strongly related to protein folding and aggregation in mammalian cells

To orthogonally validate our massive scale protein abundance measurements, we assessed the intracellular folding stability of 25 SOD1 variants, including ALS variants and aPCA variants, using a Förster resonance energy transfer (FRET) sensor measured via fluorescence lifetime imaging microscopy (FLIM) in live HEK293T cells. The sensor fuses AcGFP & mCherry to the N-and C-termini of full-length SOD1, respectively, such that stably folded SOD1 will bring the fluorescent proteins into close proximity, which decreases AcGFP fluorescent lifetime (Suppl. Fig. 3A)^47^.

We first validated that FRET occurred between AcGFP and mCherry using constructs with varied flexible linker lengths between the two fluorophores (Suppl. Fig. 3C and D). We then assayed the SOD1 variants, finding that median fluorescent lifetimes ranged from 1.76 to 2.22 ns across the variants, where SOD1-WT is 1.78 ns. All variants which are defined as stable via the aPCA assays (A4G, K30F, H48T, N53L, H63V, D90G, and S98D) with the exception of A4G, result in WT-like (1.78 ns) fluorescent lifetimes, ranging from 1.76 to 1.93 ns (Fig. 2A). Unstable variants from aPCA (I17G, L38E, N65F, R79I and V97Q) all have fluorescent lifetimes higher than SOD1-WT, ranging from 2.05 to 2.22 ns (Fig. 2A). Meanwhile, ALS-associated variants, which are found to be unstable via aPCA, have fluorescent lifetimes ranging from 2.03 to 2.19 ns (Fig. 2A and B). Conversion of fluorescent lifetimes to folding scores relative to SOD1-WT (whereby a negative score indicates decreased stability and a positive score indicates increased stability) allowed us to correlate the folding sensor outputs with aPCA, resulting in a strong anti-correlation between the two techniques (Pearson’s R=-0.78, p=4.62e-06). This suggests that mutational effects on abundance are very likely to be occurring from protein destabilization, and that these changes are mostly conserved across cellular systems (Fig. 2B).

**Figure 2.**
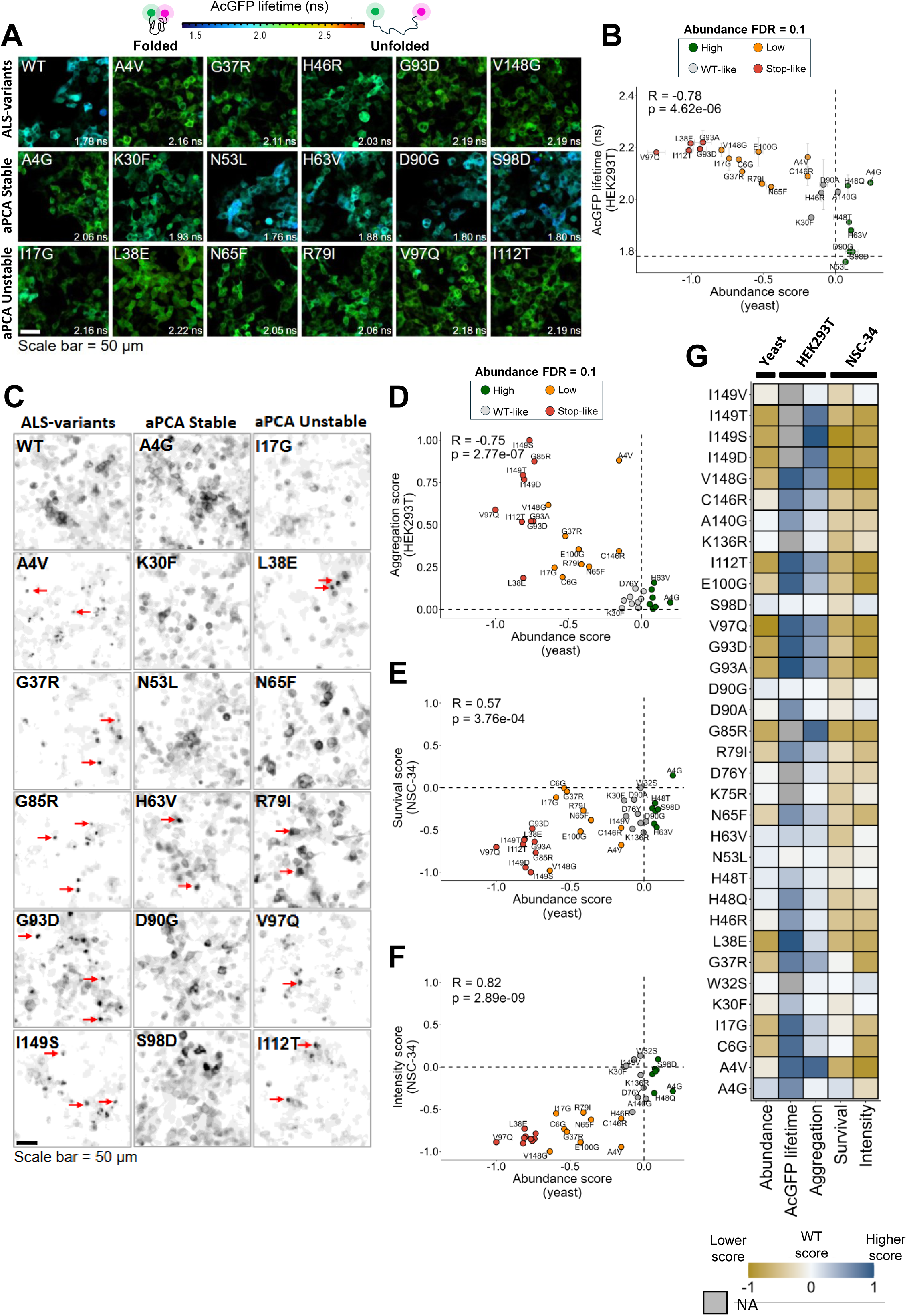
SOD1 abundance PCA in yeast is highly correlated with SOD1 variant folding, toxicity, and aggregation in mammalian cell models. **(A)** Fluorescence lifetime imaging microscopy (FLIM) of SOD1 variant folding sensors scaled according to lifetimes with median lifetime reported in nanoseconds. **(B)** Correlation between FLIM and aPCA of SOD1 variants. Pearson’s correlation coefficient and p-value are indicated. **(C)** Images of EGFP-tagged variants of SOD1 showing the formation of aggregates (bright dense clumps) of ALS variants and some unstable variants. Examples of aggregates are indicated with red arrows. Scatter plots between yeast abundance and **(D)** aggregation, **(E)** cell survival, and **(F)** fluorescence intensity. **(G)** Heatmap showing scores of all cell measurements, including yeast abundance, HEK293T fluorescence life-time (FLIM) and aggregation, and NSC-34 survival and fluorescence intensity. Scores are centered to the relative WT score and normalized by dividing each score by the maximum absolute value for each assay, resulting in scores scaling from -1 to 1.

Next, we asked whether aPCA from yeast was also correlated to other SOD1-ALS disease-relevant parameters, including protein aggregation, cell survival, and protein abundance in mammalian cells. To do so, we expressed 34 EGFP-labelled SOD1 variants including ALS-variants and variants from the DMS library in either HEK293T cells (aggregation) or NSC-34 (motor neuron-like) cells (fluorescence intensity and survival) (Fig. 2C and Suppl. Fig. 3E). Our aPCA measurements show strong quantitative relationships with all three phenotypes (Fig. 2D, E and F), however the strongest correlation is with the fluorescence intensity measurements (Pearson’s R=0.82, p=2.74e-09) (Fig. 2F). This provides strong support that mutational effects on SOD1 stability and degradation are conserved across cellular systems. More importantly, our aPCA measurements show a strong anti-correlation with aggregation propensity (Pearson’s R=-0.75, p=2.72e-07) (Fig. 2D). Variants with reduced, stop-like abundance more readily form intracellular aggregates (Fig. 2C and D), consistent with SOD1 destabilization upon mutation leading to aggregation. Finally, yeast abundance shows a moderate but significant correlation with cell survival (Pearson’s R=0.57, p=3.77e-04) (Fig. 2E). Together, these results indicate that abundance measurements provide a quantitative measurement of mutation-induced protein destabilization (Fig. 2G) that is predictive of multiple disease-relevant phenotypes across cellular systems.

Our aPCA scores also correlate with those from a previous large scale assay which employed a fluorescence-based read-out of SOD1 monomer abundance, VAMP-seq, in mammalian cells^35^ (Spearman’s rho=0.63) (Suppl. Fig. 4A). We observe a higher correlation between the two data sets in β-strands than in loops (Suppl. Fig. 4B and D) and similar effects in Greek-key loops III and VI (Spearman’s rho=0.78 and 0.90, respectively) (Suppl. Fig. 4C and D). The yeast abundance data more clearly captures the detrimental effect at zinc-binding positions, with most mutations being destabilizing in aPCA, but tolerated in VAMP-seq (Suppl. Fig. 4D), suggesting an increased sensitivity of the yeast abundance assay for mutations in these regions. Importantly, the yeast dataset has ∼2-fold more mutations than the VAMP-seq dataset (Suppl. Fig. 4E), with the addition of amino acid indels, allowing us to provide information on SOD1 backbone perturbations. By comparing the VAMP-seq dataset to our range of validation measurements in NSC-34 and HEK293T cells, we observe overall similar correlations to those observed for aPCA (Pearson’s R=-0.77, p=2.65e-07 for aggregation; R=0.55, p=9.86e-04 for cell survival; and R=0.80, p=4.44e-08 for protein abundance) (Suppl. Fig. 4G-I). However, when it comes to the FLIM data we obtain a substantially higher correlation between aPCA vs FLIM (Pearson’s R=0.78, p=4.623-06) than VAMP-seq vs FLIM (Pearson’s R=0.67, p=4.38e-04), even though both VAMP-seq and FLIM assays are carried out in the same mammalian cell line (HEK293T) (Suppl. Fig. 4F), suggesting the aPCA assay and FLIM dataset more accurately reflect the impact of mutations on SOD1 folding. This improved agreement may partially arise from the use of triplicate measurements, which enable replicate-derived uncertainty estimates ensuring reusability and providing a quantitative resource for downstream modelling and benchmarking.

### Mapping the effect of mutations on SOD1 heterodimer formation

Mutated SOD1 predominantly causes ALS in an autosomal dominant manner, meaning that WT protein is present in disease. Some evidence has suggested that heterodimers of WT SOD1 with SOD1 variants are a possible cause of toxicity in SOD1-ALS, however the number of variants assessed has been minimal. With the aim of understanding how mutations affect SOD1 heterodimerization, we performed a second deep mutational scanning assay (Fig. 1A, see Methods). Here, we measured the ability of SOD1 monomers to form heterodimers using a binding protein fragment complementation assay, Binding PCA (bPCA), also based on DHFR complementation in the presence of methotrexate (MTX). In this assay, mutant SOD1 variants are fused to one DHFR fragment, while WT SOD1 is fused to the complementary DHFR fragment (Fig. 3A). DHFR complementation now depends on the ability of SOD1 monomers to form heterodimers (Fig. 3A). bPCA replicates are highly correlated (median of Pearson’s R=0.96) (Fig. 3B and Suppl. Fig. 1D) and we obtained confident heterodimerization scores for 5939 variants including single amino acid substitutions (n=2898), single amino acid insertions (n=2758), single amino acid deletions (n=153), synonymous (n=88) and premature stops (n=42) variants (Fig. 3C).

**Figure 3.**
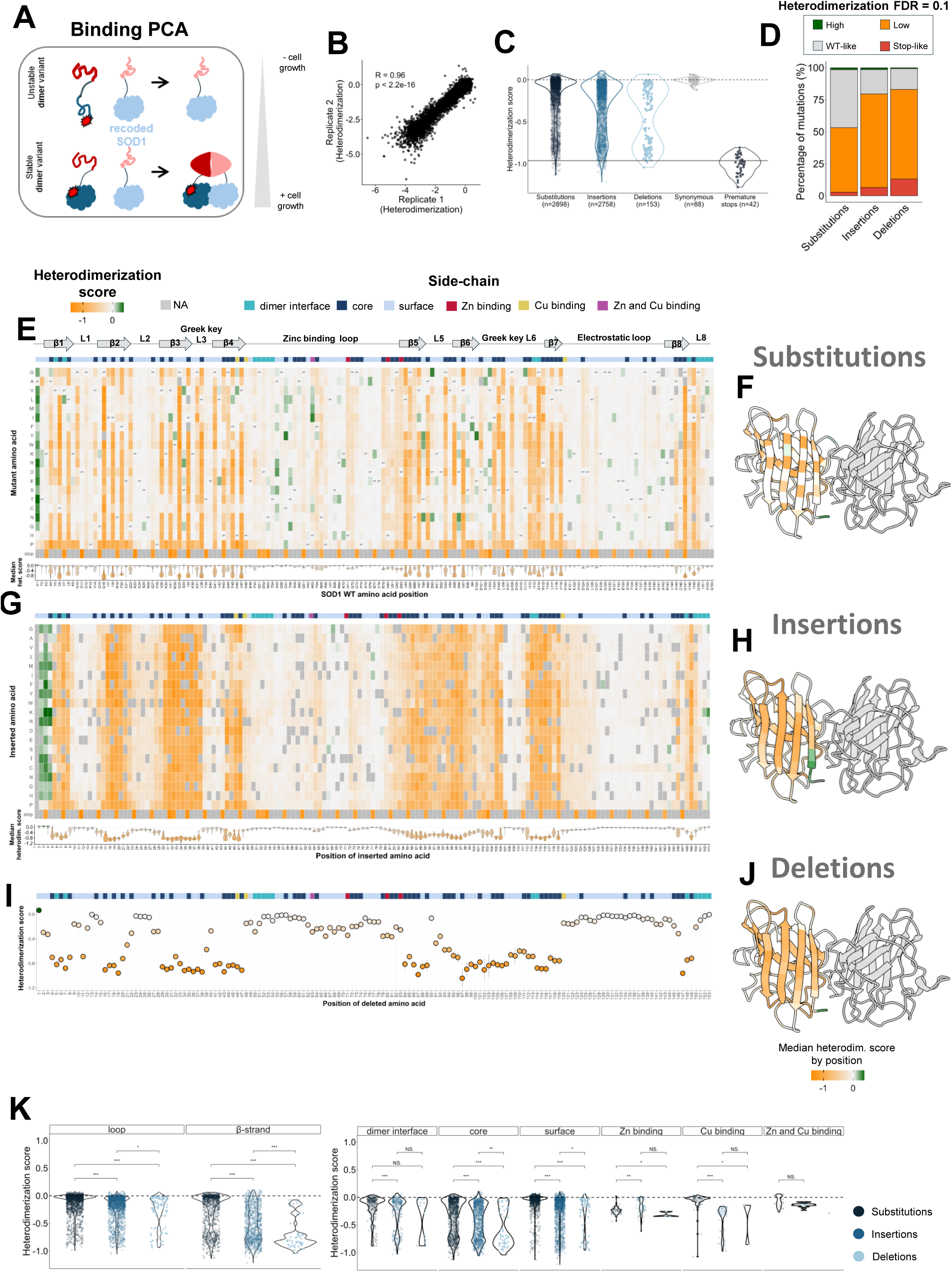
BindingPCA of SOD1 variants shows that abundance is a major determinant of heterodimerization. **(A)** BindingPCA (bPCA) workflow. Each SOD1 variant is fused to a fragment of DHFR (DHF), and a SOD1 WT copy is fused to the remaining DHFR fragment (THF). Both SOD1 WT and mutant are equally expressed **(B)** Scatter plot of bPCA replicates 1 and 2. Pearson’s coefficient and p-value are indicated. **(C)** Normalized score distribution of each variant type of the library for bPCA. The dashed line represents the WT-like score and the solid line represents the stop mutations mode. **(D)** Percentage of variants of each heterodimerization FDR category for amino acid substitutions, insertions, and deletions. Heterodimerization mutational maps for **(E)** amino acid substitutions, **(G)** insertions, and **(I)** deletions. Orange: decreased heterodimerization; grey: WT-like heterodimerization; green: increased heterodimerization. The bar over each map represents the side-chain orientation of each position (cyan: dimer-interface residues; darkblue: core residues, rSASA dimer < 25; lightblue: surface-exposed residues (rSASA dimer > 25); red: zinc-binding residues; purple: copper-binding residues. SOD1 3D dimer structures (PDB: 2V0A) colored by the median heterodimerization scores per positions of substitutions **(F**), insertions **(H**), and deletions **(J).** (**K)** Violin plots showing the heterodimerization scores distribution of mutations in loops, β-strands, and according to the side-chain.Significance is calculated with the Mann-Whitney U test: p>0.05 (NS, non-significant), p<0.05 (*), p<0.01(**), p<0.001(***).

The dynamic range of bPCA also clearly separates scores for synonymous and premature stop mutations (Fig. 3C). Overall, we observe a minor proportion of WT-like mutations with respect to the abundance measurements in the three types of variants (45% of substitutions, 19% of insertions and 16% of deletions) (Fig. 3D). More importantly, low and stop-like heterodimerization variants predominate over the WT-like heterodimerization variants (Fig. 3D).

Similar to the aPCA, substitutions result in alternating effects in β-strands, with surface exposed residues better tolerating variants than those that are buried (Fig. 3E and F), supporting that SOD1 dimers form through relatively stable conformers. Indels are instead detrimental across the entire β-strands, regardless of side-chain orientation, consistent with backbone mutations (i.e., indels) having more severe effects on protein stability than substitutions, as previously shown by Topolska *et al.*^44^. (Fig. 3G-J). Loops better tolerate variants than β-strands for all three types of variants (Fig. 3K). However, there are different levels of tolerance when comparing substitutions with indels. Core, surface-exposed and zinc-binding residues show different levels of tolerance according to mutation types, with deletions having the most detrimental effect (Fig. 3K). When it comes to dimer interface residues, mutational effects distribute bimodally, with a higher detrimental impact from indels (Fig. 3K).

### Abundance predicts heterodimerization with interface-specific residual effects

When comparing mutational effects on abundance and heterodimer formation, we find a high correlation between scores (Spearman’s rho=0.92, p<2.2e-16), revealing that the two phenotypes are strongly coupled (Fig. 4A). To model the relationship between abundance and heterodimerization, we fitted a locally weighted regression (LOWESS) curve, allowing us to capture the non-linear relationship between the two phenotypes (Methods). LOWESS fits are useful to accurately determine residuals, with residuals from the fit (from now heterodimerization (HD) residuals) defined as the difference between the observed heterodimerization scores and the heterodimerization scores expected based on abundance (Fig. 4A). We use the HD residuals to analyze the dependence of heterodimerization on protein abundance.

**Figure 4.**
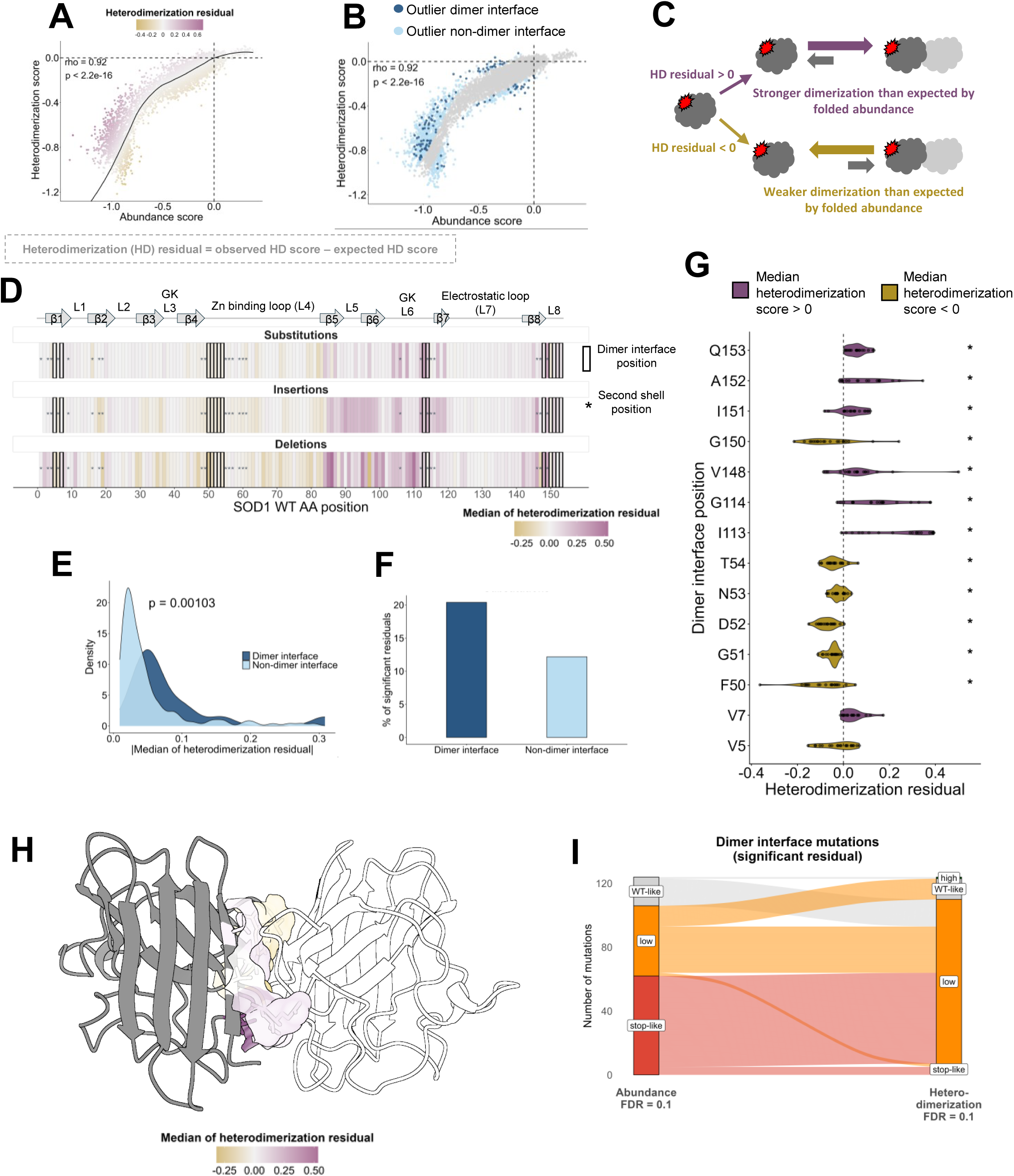
Dissecting mutational effects on protein abundance and heterodimerization. **(A)** Correlation of normalized abundance (x-axis) and heterodimerization (y-axis) scores. Dots are coloured according to the residual of the LOESS model (violet: residual >0, grey: residual ∼ 0, gold: residual < 0). The heterodimerization residual (HD) is calculated as the difference between the observed heterodimerization score and the expected heterodimerization score based on abundance by the model. Black line represents the fitting model; dashed lines represent WT scores. Spearman’s correlation coefficient and p-value are indicated. **(B)** Correlation of normalized abundance and heterodimerization scores coloured by outliers. Mutations are considered outliers when their residual is > 2*MAD (Median Absolute Deviation). Darkblue: outlier mutations of the dimer interface; Lightblue: outlier mutations out of the dimer interface. Spearman’s correlation coefficient and p-value are indicated. **(C)** Schematic representation of the heterodimerization residual effect at the molecular level. Mutations with a positive HD residual (violet) feature a stronger heterodimerization than what is expected by their abundance. Mutations with a negative HD residual (gold) a weaker heterodimerization than what is expected by their abundance. **(D)** Median heterodimerization residual by position heatmap for substitutions, insertions and deletions. Dimer interface positions are indicated with a black square, and second shell positions with an asterisk (*). **(E)** Comparison of the intensity of HD residual in dimer (darkblue) and non-dimer interface residues (lightblue). The x-axis represents the modulus of the median HD residual by position. The p-value is indicated. **(F)** Comparison of the percentage of significant HD residuals (>2MAD) in dimer and non-dimer interface residues. Significance is calculated with the Mann-Whitney U test. **(G)** Distribution of HD residuals in dimer interface residues. Violins are coloured according to whether the median HD residual at a position is higher (violet) or lower (gold) than 0. Asterisks indicate positions with a significant deviation from 0 (two-sided one-sample Wilcoxon signed-rank test, Benjamini–Hochberg adjusted p-value < 0.05). **(H)** SOD1 3D structure (PDB: 2V0A) where dimer interface positions are coloured by their median HD residual. **(I)** Alluvial plot representing changes in FDR classifications between Abundance (left) and Heterodimerization (right).

Mutations with positive HD residuals have higher scores on heterodimerization relative to their folded abundance, and mutations with negative HD residuals feature weaker heterodimerization relative to their folded abundance (Fig. 4A-C). Negative HD residuals are enriched for mutations at the SOD1 N-terminus, while positive HD residuals are enriched for mutations at the C-terminus (Fig. 4D and Suppl. Fig. 5A-F).

Dimer interface residues are enriched (1.67 fold) with significant HD residuals (residual>2* Median Absolute Deviation (MAD)) compared to non-interface positions (Mann-Whitney U test, p=1.03e-03) (Fig. 4E and F). We see a similar enrichment when including indels in the residual analysis (Mann-Whitney U test, p=6.32e-03) (Suppl. Fig. 5G and H). At the dimer interface, the enrichment of significant HD residuals is driven by positive HD residuals from mutations at positions V7, I113, G114, V148, I151, A152 and Q153, suggesting that monomer destabilizing mutations at these positions may get partially stabilized in the dimer context (Fig. 4G and H and Suppl. Fig. 5I). Negative HD residuals are instead enriched at positions F50, G51, D52, N53, T54 and G150 (Fig. 4G and H and Suppl. Fig. 5I) and the same positional enrichment is reflected in indels (Suppl. Fig. 5J and K). Interestingly, the deletion of position F50 features as the highest negative HD residual, consistent with previous work^12^ suggesting a crucial role of this position for SOD1 dimerization (Suppl. Fig. 5K). Dimer interface substitutions and insertions are 3 fold more enriched with positive HD residuals than with negative ones, while deletions have 1.85 fold more negative HD residuals than positive ones, suggesting that deletions of specific interface residues mainly compromise SOD1 dimerization (Suppl. Fig. 5I).

Finally, when considering only mutations with significant HD residual at the interface, we observe a wide range of mutational effects in both phenotypes, including for example mutations that only affect dimerization (high/WT-like abundance and low heterodimerization, FDR=0.1), mutations that only affect abundance (low abundance and WT-like heterodimerization, FDR=0.1) and mutations that affect dimerization more than abundance (low abundance and stop-like heterodimerization, FDR=0.1) (Fig. 4I).

### Identification of allosteric mutations modulating SOD1 dimerization

As expected, many mutations at the dimer interface feature larger HD residuals compared to non-interface positions (Mann-Whitney U test, p=1.03e-03) (Fig. 4E and F). However, mutations outside the dimer interface can occasionally have large HD residuals. We propose these mutations impact heterodimer formation via allosteric-like mechanisms. Among them we can distinguish two groups: with positive or negative effects on dimerization (from now on, positive or negative allosteric mutations, respectively). We identify 661 allosteric substitutions (418 positive and 243 negative HD residuals (|HD residual|>|median interface HD residual|)) distributed along the SOD1 structure (Fig. 5A). Residues in direct contact with interface residues (second-shell residues) have 1.3 fold more allosteric variants than other allosteric residues located far from the interface (Chi-square test, p=1.39e-07) (Suppl. Fig. 6A).

**Figure 5.**
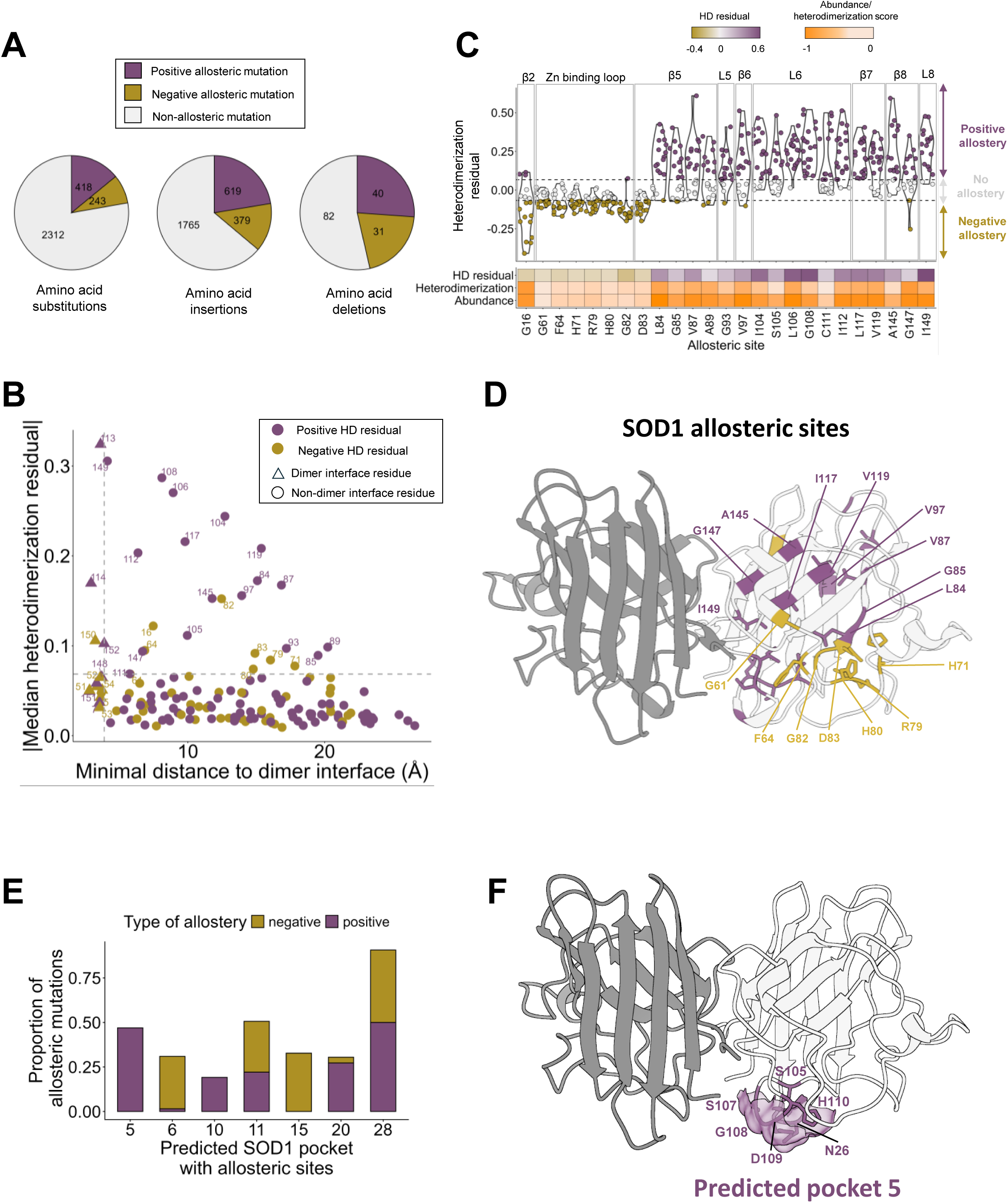
SOD1 dimerization can be allosterically modulated. **(A)** Pie chart of allosteric and non-allosteric substitutions, insertions and deletions. **(B)** Relationship between per-position median HD residual to the minimum atom distance to the dimer interface. Horizontal dashed line: absolute HD residual median of dimer interface residues; Vertical dashed line: highest minimal distance of dimer interface residues. **(C)** Distribution of the HD residuals of mutations in allosteric sites. Top dashed line: positive median HD residual of dimer interface residues; Bottom dashed line: negative median HD residual of dimer interface residues. **(D)** Mapping of allosteric sites in the SOD1 3D structure (PDB: 2V0A). **(E)** Proportion of allosteric mutations found in the predicted SOD1 pockets with allosteric sites. **(F)** SOD1 3D structure (PDB: 2V0A) with residues forming the predicted pocket 5 represented in violet as sticks and surface.

By plotting the HD residuals of mutations/residues against the minimal distance to the dimer interface in the SOD1 3D structure, our results reveal an allosteric decay on the effect of mutations on heterodimerization: mutations at interface residues have the highest HD residuals followed by second-shell residues, with weaker allosteric effects as the distance to the dimer interface increases (Fig. 5B and Suppl. Fig. 6B). The same plot identifies allosteric sites, defined as positions where their median HD residual of mutations is higher than the median HD residual of dimer interface residues (Fig. 5B) (Methods). We identify 25 allosteric positions distributed through β-strands 2, 5, 6, 7 and 8, the zinc binding loop and loops 5 and 6, where the enrichment of allosteric mutations is at least 2-fold with respect to non-allosteric positions, with 17 positions featuring positive allostery and 8 with negative allostery [odds ratio (OR)>2, FDR<0.1, Fisher’s Exact Test (FET)] (Fig. 5B-D, Suppl. Fig. 6C). Of these 23 allosteric sites, 6 are second shell positions. However, only at three positions (G82, G108 and I117) all substitutions (19/19) have an allosteric effect: at position G82, all but one substitution are negative allosteric mutations, whereas all substitutions at positions G108 and I117 are positive allosteric mutations (Fig. 5C). Positions enriched with positive allosteric mutations are overall more frequent (OR=12.54, FDR<0.1 and OR=6.58, FDR<0.1, FET, for negative allosteric mutations) (Suppl. Fig. 6D).

Next, we analyzed how the allosteric signal is potentially propagated. We hypothesize that the allosteric signal can be propagated between two positions in physical contact (distance between any atom<5Å) and with the same allosteric effect. We observe that the negative allosteric signal is propagated in two paths: from G82 to the second shell residue F64 and through a second path that includes residues H71, R79, D83, H80, G82 and F64 (Suppl. Fig. 6E and F). The positive allosteric signal propagation may end in different second shell positions: L106, I112, I117, G147 or 108 (Suppl. Fig. 6G and H).

We next asked whether our identified allosteric sites may be therapeutically targetable. We predicted putative ligandable pockets in SOD1 (Methods) and examined whether they contain allosteric residues (Suppl. Fig. 7A). Of the 15 non-interface predicted pockets, we identify 7 containing allosteric residues (Fig. 5E and Suppl. Fig. 7A). Notably, pocket 5 and, to a lesser extent, pocket 10 are enriched exclusively in positive allosteric mutations. Since mutations in pocket 5 show stronger dimerization relative to abundance, we speculate that perturbations in the same pocket can potentially have dimerization-favouring effects. We also note that pocket 15 is exclusively enriched with negative allosteric mutations. Pockets 6, 11, 20, and 28 instead feature a mixture of both classes (Fig. 5E and F, Suppl. Fig. 7A-F). These findings suggest that SOD1 dimerization may be modulated through multiple structurally distinct pockets and point at pocket 5 as an accessible site through which small molecules may allosterically enhance dimerization.

### Amino acid insertions and deletions have stronger allosteric effects than substitutions

Our SOD1 DMS atlas also contains information about the effect of deletions and insertions in SOD1 on abundance and heterodimerization, allowing us to probe allostery due to not only side-chain but also to backbone perturbations. We find that 36% (n=998) of insertions and 46% (n=71) of deletions have an allosteric-like effect on SOD1 heterodimerization (Fig. 5A), compared to 22% allosteric substitutions. In both insertions and deletions, as in substitutions, the predominant allosteric effect is positive (∼21% of insertions and ∼25% of deletions), with a smaller proportion of indels having negative allosteric effects (∼12% of insertions and ∼17% of deletions) (Fig. 5A). Negative allosteric mutations are mainly located at the N-terminus of SOD1 for both insertions and deletions (Suppl. Fig. 5C-F), with a higher enrichment of allosteric insertions in β-strands 2 and 3 (Suppl. Fig. 5C and D).

When comparing allosteric effects across substitutions, insertions and deletions, we observe a moderate correlation of residuals (Suppl. Fig. 8A-C), with the highest correlation being for insertions vs deletions (Pearson’ R=0.54, p=1e-12) (Suppl. Fig. 8C). When comparing position-specific allosteric effects across mutational classes, we find that the direction of allostery (whether it is positive or negative) is generally conserved, with the same allosteric effect predominating across mutation types (Suppl. Fig. 8D–F). However, substitutions and indels occasionally produced opposite allosteric effects (Suppl. Fig. 8D–F), suggesting specific allosteric mechanisms for indels backbone perturbations.

### The majority of pathogenic variants reduce protein abundance and dimerization

Accurate interpretation of genetic variants is essential for understanding disease mechanisms and for determining whether newly identified variants contribute to disease. Clinical classification frameworks increasingly incorporate evidence from functional assays to support pathogenicity assessments. Using our two comprehensive variant effect maps, we therefore assessed if monomer abundance and heterodimerization can contribute to clinical variant interpretation for SOD1-ALS.

Currently, 112 missense and 1 inframe deletion mutations in SOD1 are reported as pathogenic or likely pathogenic (from now on P/LP), 54 missense and 1 inframe deletion as VUS and 16 variants have conflicting classifications of pathogenicity (ClinVar data from Feb2026). Our dataset contains scores for all pathogenic variants except three. We observe that P/LP missense variants distribute bimodally in abundance (Fig. 6A), with 74 (68%) classified as low/stop-like abundance and 35 (26%) as WT-like abundance (Fig. 6B (i); see Supplementary Dataset S6). When it comes to heterodimerization, 25 P/LP variants are classified as WT-like (23%), 23 of which are also WT-like in abundance (Fig. 6A and B (i)). However, 2 P/LP variants (C146R and I149T) feature low abundance but WT-like dimerization, suggesting that the destabilization of monomeric SOD1 may get partially compensated by heterodimer formation (Suppl. Fig. 9A). None of the 112 P/LP variants occur at the dimer interface, with two P/LP variants featuring a stronger heterodimerization than was expected based on their abundance (Fig. 6B (ii)). On the other hand, 24/112 variants occur in second shell residues, with four variants having weaker heterodimerization relative to their abundance (Fig. 6B (ii)). Three of these four are negative allosteric variants, G16S, G16C and H71T (Fig. 6C), suggesting their pathogenic outcome could involve an allosteric mechanism that impairs dimer formation.

**Figure 6.**
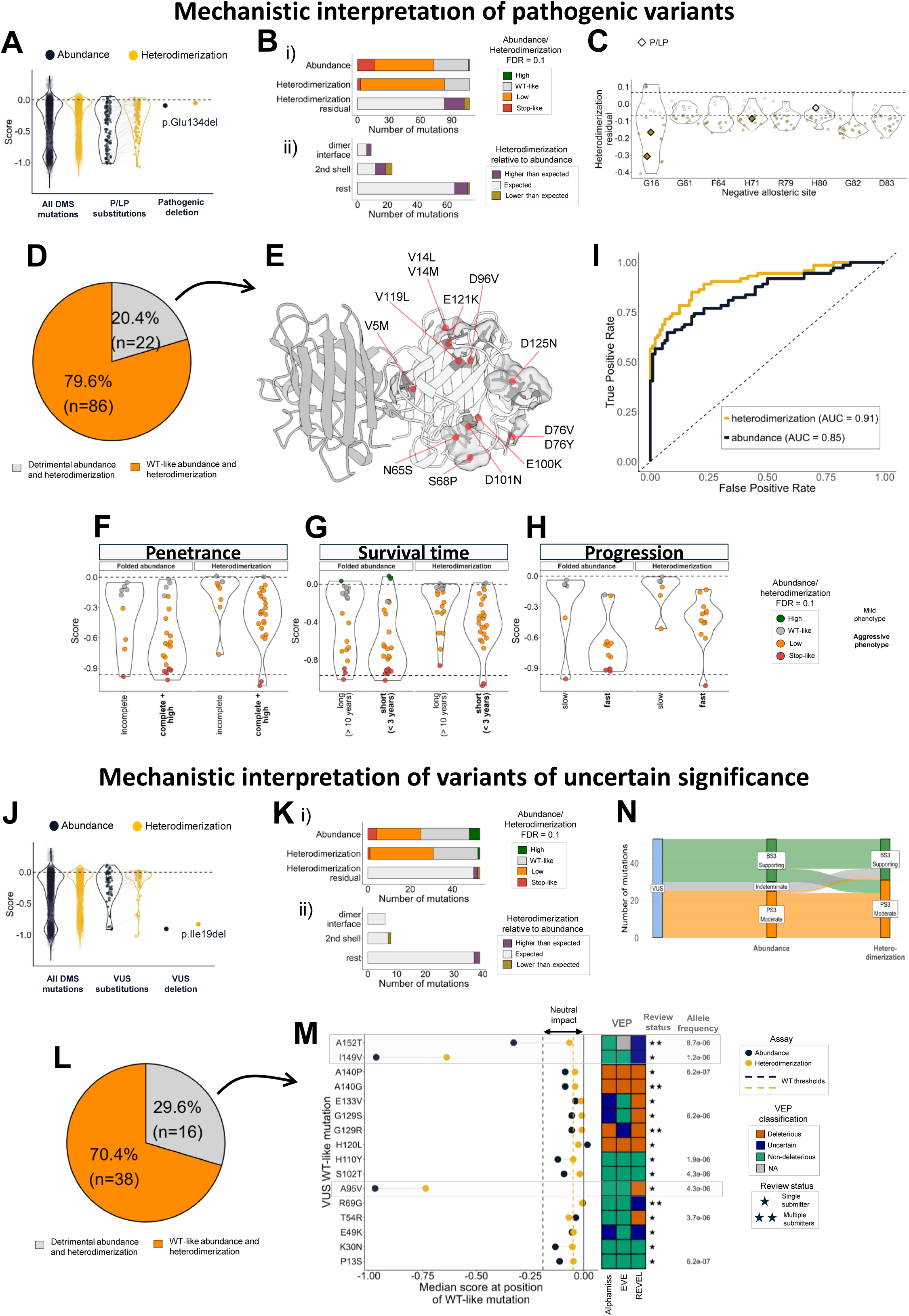
SOD1 multiphenotyping for clinical variant interpretation. **(A)** Violin plot showing the abundance/heterodimerization scores of all the mutations measured in this work (substitutions and indels), and the reported pathogenic (P)/likely pathogenic (LP) variants from gnomAD and ClinVar. **(B)** Number of P/LP variants within each FDR category based on abundance, heterodimerization, and HD residuals. **(C)** Distribution of HD residuals of negative allosteric sites, highlighting the P/LP variants as diamonds. Top dashed line: positive median HD residual of dimer interface residues; Bottom dashed line: negative median HD residual of dimer interface residues. **(D)** Pie chart showing the proportion of P/LP variants with low/stop-like classification (Orange) and WT-like classification (grey) in both abundance and heterodimerization. **(E)** Mapping of P/LP WT-like variants in the SOD1 3D structure (PDB: 2V0A). Distribution of abundance and heterodimerization scores for disease penetrance **(F),** survival time **(G)** and disease progression **(H)**. **(I)** Receiving Operator Characteristic (ROC) curve showing the performance of abundance and heterodimerization at classifying pathogenicity. The dataset used in this ROC is composed of 74 P/LP variants and 96 “benign” controls (including 86 synonymous, 1 likely benign and 9 relatively frequent variants). **(J)** Violin plot showing the abundance/heterodimerization scores of all the mutations measured in this work (substitutions and indels), and the reported VUS. **(K)** Number of VUS variants with each FDR category based on abundance, heterodimerization, and HD residuals categories. **(L)** Proportion of VUS with low/stop-like classification (orange) and WT-like classification (grey) in both abundance and heterodimerization. (**M)** Comparison of VUS features relevant for clinical variant interpretation. From left to right: comparison of each VUS WT-like (n=16) with the median abundance/heterodimerization score at the position they occur. VUS that occur at positions where the median of scores is lower than the neutral impact range are highlighted in grey; the VEP column indicates the classification of Alphamissense, EVE and REVEL; the review status column indicates the strength of evidence for each WT-like VUS based on the number of submitters. **(N)** Alluvial plot showing the BS3/PS3 evidence of all VUS in both abundance and heterodimerization.

Overall, our dataset assigns 79.6% (n=86) of P/LP variants as destabilizers of both the SOD1 monomer and the dimer (Fig. 6D), consistent with protein destabilization as the dominant mechanism driving SOD1-ALS. This is also in agreement with previous work demonstrating that protein destabilization is the main but not the only mechanism driving disease ^48^. Overall, the impact of P/LP variants on dimer formation correlates with that in abundance (Spearman’s rho R=0.94, p<2.2e-16) (Suppl. Fig. 9B). However, among these destabilizing variants, 24 have a significant residual, suggesting they have a distinct detrimental impact on dimerization and abundance (Fig. 6B).

### Pathogenic variants not impairing protein abundance and dimerization cluster away from the dimer interface

Although 79.6% of P/LP variants have a destabilizing effect on both abundance and heterodimerization, there are still 20.4% of P/LP variants (n=22), including the pathogenic deletion E133del, that have no significant effect on either phenotype (z-test, abundance FDR=0.1, heterodimerization FDR=0.1) (Fig. 6A and D). These pathogenic variants, labeled as WT-like on the basis of dual phenotyping, likely cause disease via mechanisms other than destabilization and dimer impairment. Mapping these variants on the SOD1 3D structure reveals they cluster in relatively high solvent-accessible regions that are distant from the dimer interface (Fig. 6E), including positions at the zinc-binding loop, the electrostatic loop and exposed residues of β-strands 5, 6, 7 and 8 (Suppl. Fig. 9C).

We investigated these 22 variants further by comparing our DMS datasets with information available in public databases. First, we analyzed the median mutational effect on abundance and heterodimerization at the positions where these mutations occur. 9/22 variants occur in positions where the median abundance/heterodimerization impact is detrimental (V5M, V14L, V14M, A89V, V119L and V148I) (z-test; median abundance score<-0.19, FDR=0.1; median heterodimerization score<-0.05, FDR=0.1) (Suppl. Fig. 9C). The remaining mutations occur in positions where the median mutational impact is neutral (Suppl. Fig. 9C). We also observe that many of these P/LP variants are assigned as non-deleterious or indeterminate scores by Variant Effect Predictors (VEPs). Most of these P/LP WT-like variants occur in loops and surface residues (Suppl. Fig. 9D), especially in the zinc and electrostatic loops (Suppl. Fig. 9C-E) where mutations are better tolerated than in other regions, suggesting they cause disease via mechanisms or interactions not explored in this work.

### Variants impairing abundance and dimerization have more severe clinical phenotypes

By relating our data to published ALS datasets reporting variant penetrance, survival time and progression^1,26,49–53^ (see Methods), we observe variants with a broad range of effects on both abundance and heterodimerization in virtually all categories (Fig. 6F and H). Overall, mutations defined as WT-like on the basis of their impact on abundance or heterodimerization are significantly enriched in the milder phenotypes (i.e., incomplete penetrance, long survival time and slow progression) (FET, all p-values<0.05). However, two variants (V7E and H48Q) feature higher abundance than WT and have been reported in cases of ALS with short survival time (z-test; abundance score>-0.01, FDR=0.1). Notably, H48Q combines high abundance with low heterodimerization (z-test; abundance score>-0.01, heterodimerization<-0.05, FDR=0.1) (Fig. 6G).

We also note that variants associated with more aggressive phenotypes (i.e., complete penetrance, short survival time and fast progression) have larger residuals than those associated with less aggressive phenotypes (incomplete penetrance, long survival time and slow progression) (Suppl. Fig. 9F). This suggests that clinical aggressiveness may not be explained by reduced monomer abundance alone, but may also involve abundance-independent defects in heterodimerization. Notably, variants with large negative residuals are observed exclusively among aggressive phenotypes, consistent with a disproportionately strong impairment of heterodimerization beyond what would be expected from their effects on abundance (Suppl. Fig. 9F).

### Reduced abundance and dimerization contributes moderate evidence towards pathogenicity

We next set out to use our abundance and heterodimerization scores to define likelihood ratios (LR) of pathogenicity. We further curated SOD1 missense variants from public databases (ClinVar, ALSoD, LOVD) using modified American College of Medical Genetics and Genomics and the Association for Molecular Pathology (ACMG/AMP) classification criteria^54^. To avoid circularity, we explicitly excluded evidence arising from functional assays when classifying the variants. The result is a calibration set of 74 variants classified as LP/P and only a single variant classified as likely benign (N19S; see Supplementary Dataset S7). When lacking well-classified benign controls in a gene, the Genome Aggregation Database (https://gnomad.broadinstitute.org/)^55^ can be used as a source of common variation^54^. However, for SOD1 the most common missense variant is an established pathogenic variant (D90A) and many of the other missense variants are too rare to be used as “benign” controls given the prevalence and adult age-of-onset of ALS. We therefore used 86 synonymous variants from our dataset, the likely benign variant N19S, and only 9 unclassified (not present in ClinVar) relatively common missense variants (>5 allele counts in gnomAD) whose frequency in extensively sequenced population cohorts makes a highly penetrant pathogenic effect unlikely. We calculated LRs for pathogenicity using 74 LP/P variants and 96 “benign” control variants and converted them to ACMG/AMP PS3/BS3 evidence strengths, with abnormal function defined as stop-like/low function and normal function as WT-like. By applying these criteria, using each dataset, we can assign pathogenic moderate evidence (PS3 Moderate) for variants with stop-like or low abundance/heterodimerization and benign supporting evidence (BS3 Supporting) for variants with WT-like abundance/heterodimerization.

122 out of 149 variants that have indeterminate evidence based on abundance can be assigned either BS3 Supporting (n=119) or PS3 Moderate (n=3) based on heterodimerization (Suppl. Fig. 9E), demonstrating that multidimensional phenotyping can improve clinical variant interpretation and our understanding on how SOD1 variants contribute to disease.

We used the same reference set to evaluate the performance of abundance and heterodimerization as classifiers of SOD1 variant deleteriousness. Receiver operation characteristic (ROC) curves show that heterodimerization performs better than abundance (heterodimerization AUC=0.91, abundance AUC=0.85; Delong test, Z=2.97, p=0.003) (Fig. 6I). We also asked whether a combined score that integrates both abundance and heterodimerization data would perform better than individual scores. We defined different combined scores by assigning different weights to each phenotype, but regardless of the weights, a combined score performs equally to heterodimerization alone (Suppl. Fig. 9G). Altogether, this suggests that all the abundance information is contained in the heterodimerization phenotype and, in consequence, heterodimerization provides additional mechanistic information that cannot be explained entirely by abundance.

### The majority of VUS have a detrimental impact on abundance and heterodimerization

We next applied this mechanistic information to variants of uncertain significance, variants for which clinical interpretation remains unresolved.

The abundance scores distribution of VUS is mainly unimodal, with a dominant peak of WT-like/slightly destabilizing mutations (Fig. 6J). We classify 48% of VUS (n=26) as low/stop-like abundance, 42% (n=23) as WT-like and 9% (n=5) as high abundance (Fig. 6K (i)). When it comes to heterodimerization, ∼60% (n=32) of VUS are classified as low/stop-like, and 40% (n=22) featuring WT-like heterodimerization (Fig. 6J and K (i)). As in pathogenic variants, abundance and heterodimerization of VUS are highly correlated (Spearman’s rho = 0.88, p<2.2e-16) (Suppl. Fig. 10A). Although six VUS occur at the SOD1 dimer interface, their effect on dimerization can be explained by the abundance effect (|HD residual| < |median interface HD residual|). There is one second shell VUS with a stronger impairment of dimerization than what is expected by its folded abundance, and two VUS located away from the interface region that feature a stronger dimerization relative to their folded abundance, suggesting a stability compensation upon dimerization (Fig. 6K (ii)).

More importantly, our dataset provides quantitative evidence that 70.4% of VUS destabilize both monomer folding and dimerization (low or stop-like abundance/heterodimerization) (Fig. 6L), including the deletion of residue I18 (Fig. 6J). We also identify VUS that have a neutral effect on abundance but are detrimental for dimerization (Suppl. Fig. 10B). Moreover, we find 16 VUS that are WT-like in both assays: 9 occur in positions where the median effect of mutations on abundance and dimerization is neutral (Fig. 6M) while 5 of them occur in positions where many other mutations are instead detrimental. VEP classification for these variants is mixed (Fig. 6M).

According to our calibration framework, we can assign PS3 Moderate evidence for those 70.4% of VUS that are destabilizing. In contrast, the evidence for the remaining 16 WT-like VUS is BS3 Supporting (Fig. 6L). Caution should be applied though as variants with WT-like abundance and heterodimerization can also be pathogenic. Thus, the neutral effect of these VUS is still compatible with these variants causing disease by other mechanisms beyond destabilization. While 48/54 VUS variants can be assigned either BS3 Supporting or PS3 Moderate based simply on abundance, the remaining 6 which have indeterminate evidence based on abundance, can be classified as BS3 Supporting based on heterodimerization (Fig. 6N). In summary, heterodimerization data could contribute evidence to 100% of current SOD1 VUS.

Finally, we also investigated SOD1 variants that are not well classified due to conflicting interpretations and found that 9/16 of these variants are detrimental for both phenotypes, while 5/16 have a neutral impact (Suppl. Fig. 10C-E). Abundance and heterodimerization of these variants strongly correlate (Spearman’s rho = 0.92, p<2.2e-16) (Suppl. Fig. 10F). Of the 16 variants with conflicting classifications of pathogenicity, we propose BS3 Supporting evidence for 9 and PS3 Moderate evidence for 7 in abundance (Suppl. Fig. 10G).

### Abundance and heterodimerization scores classify deleteriousness better than VEPs and previous SOD1 DMS

To evaluate the performance of abundance and heterodimerization with the performance of VEPs and the previous SOD1-VAMP-seq at classifying deleteriousness, we used the reference set that was generated by Axakova *et al.*^35^(Methods). We note that heterodimerization (AUC=0.85) still performs better that abundance (aPCA) (AUC=0.81), abundance (VAMP-seq) (AUC=0.80), and outperforms the classification performance of three VEPS, including Alphamissense (AUC=0.76), EVE (AUC=0.76) and ESM-b1 (AUC=0.73) (Suppl. Fig. 10H).

## DISCUSSION

The multidimensional DMS approach presented here provides a comprehensive systematic view of how all mutations in SOD1 perturb monomer abundance and heterodimerization, linking these molecular effects to downstream cellular phenotypes and showing utility in clinical variant interpretation. While a recent DMS study of SOD1 measured the effect of missense mutations on protein abundance^35^, this work combines dual-phenotype mapping with extensive orthogonal validation across multiple cell models and measurements. The strong quantitative agreement between yeast abundance measurements, FLIM-based folding measurements, aggregation quantification in HEK293T cells, and cell survival in motor neuron-like cells indicates that the mutational effects captured here reflect fundamental biophysical constraints on SOD1 monomer and dimer stability that are conserved across systems. Together with the scale of the dataset and its associated uncertainty estimates, these features establish a quantitative resource that is readily reusable for benchmarking computational predictors, developing mechanistic models, and integrating future data.

By profiling substitutions, insertions, and deletions, this work also expands variant effect mapping beyond the missense space. Although indels overall show stronger detrimental effects on both abundance and heterodimerization, the correlation with substitutions suggests shared mutational constraints across different mutational classes. Importantly, however, indels also reveal larger and distinct allosteric-like effects that would not have emerged from substitutions alone.

Despite a strong coupling between abundance and heterodimerization, the dimerization assay provides additional mechanistic and clinical information. Heterodimerization scores explain the deleterious effects of pathogenic variants slightly better than abundance alone. This is biologically coherent with SOD1 functioning as an obligate dimer, where native assembly integrates both monomer and dimer stability. Importantly, ∼80% of pathogenic variants and ∼70% of VUS perturb both abundance and heterodimerization, consistent with SOD1 destabilization as a major driver of ALS. Overall, our datasets can provide moderate evidence for pathogenicity and supporting evidence for benignity.

Decoupling the abundance and heterodimerization datasets reveals variants with disproportionate impacts on the dimeric state that cannot be entirely explained by monomer destabilization. These heterodimerization residual effects map to the interface but also to distal positions where mutations have allosteric effects. Identifying those allosteric sites where mutations have a positive effect on dimerization allows us to map druggable pockets in SOD1 that could represent targets for small molecule stabilizers of the native SOD1 dimer. In contrast to previous strategies that target the poorly accessible dimer interface or rely on covalent stabilization, these pockets may provide more tractable sites for therapeutic intervention.

Our data reinforces the relationship between SOD1 destabilization and aggregation in ALS. Variants with reduced abundance strongly anti-correlate with aggregation propensity in mammalian cells, consistent with destabilization increasing access to aggregation-prone states. At the same time, the existence of a small set of pathogenic variants with WT-like abundance and limited aggregation, including ∼20% of pathogenic variants (n=22) that are WT-like in both assays, suggests the existence of alternative mechanisms of pathogenesis, in line with the mechanistic heterogeneity observed clinically across SOD1 variants. These neutral variants interestingly cluster away from the dimer interface and likely drive pathogenesis through alternative pathways, such as disrupted metal coordination, which will require distinct functional assays to be resolved. This observation highlights the need for caution when using functional assays to infer benignity, as pathogenic variants may operate through mechanisms not captured by a given assay.

When applying our mechanistic information to clinical phenotype interpretation (penetrance, survival time and progression), we observe that slightly destabilizing variants are found in milder phenotypes (incomplete penetrance, long survival time and slow disease progression) but not in the more aggressive phenotypes (complete or high penetrance, short survival time and fast disease progression), a trend which is more evident in the heterodimerization dataset. This opens the door to explore the use of functional assays to inform patient stratification. The limited number of patients for which this information is available hinders our ability to make stronger conclusions. The quantification of other disease-relevant mechanisms, including the exploration of variants that act via allosteric mechanisms, can potentially provide a more solid discrimination between phenotypes^56^.

More broadly, these findings highlight the importance of multidimensional phenotyping for mechanistic variant interpretation. A single phenotype is insufficient to capture the mechanistic diversity of SOD1 mutations. Comparing multiple phenotypes not only improves variant classification, but also reveals relationships that would otherwise remain hidden. This approach identifies variants that primarily affect folding stability, variants that disproportionately perturb dimerization, and variants that appear neutral in both assays despite clinical pathogenicity. Together with the identification of accessible allosteric pockets, these results highlight how multidimensional variant effect mapping can improve mechanistic understanding, clinical variant interpretation, patient stratification, and the development of mechanism-informed therapeutic strategies for individuals carrying SOD1 variants.

## Supporting information

Description Supp Files

## Acknowledgements

Work in the lab of B.B. is supported by the La Caixa Research Foundation project ‘ATLAS-ALS’ (LCF/PR/HR24/00672), by the Spanish Ministry of Science, Innovation and Universities (PID2021-127761OB-I00, PID2024-157544OB-I00 and RYC2020-028861-I, funded by MCIN/AEI/10.13039/501100011033, “ERDF A way of making Europe” and “ESF Investing in your future”) and by the European Union (ERC Consolidator, Glam-MAP, 101125484). Views and opinions expressed are, however, those of the author(s) only and do not necessarily reflect those of the European Union or the European Research Council. Neither the European Union nor the granting authority can be held responsible for them. IBEC is a member of the CERCA Program/Generalitat de Catalunya. Work in the lab of L.M. is supported by Motor Neurone Disease Australia (Jenny Simko Innovator Grant 2023), La Caixa Research Foundation project ‘ATLAS-ALS’ (LCF/PR/HR24/00672), FightMND, and NHRMC. The authors acknowledge the facilities, and the scientific and technical assistance of the Fluorescence Analysis Facility at Molecular Horizons, University of Wollongong. We acknowledge Siobhan Suters and Jordan Cater for technical assistance.

Finally, we thank the Lehner Lab for providing the plasmids used in this study and for helpful discussions of the data and the CRG Genomics core technology for all the support with sequencing.

## Data availability

Raw sequencing reads are deposited in the European Nucleotide Archive (ENA) as part of the study PRJEB114475. The processed reads and abundance/heterodimerization scores are available as Supplementary Dataset S2.

## Author Contributions

B.B, L.M., and B.A.T. conceived the study. T.Q., L.A., L.R., D.B. and M.M. developed the methodology and performed the experiments. T.Q., L.A., L.R., D.B., B.A.T., L.M. and B.B. analyzed and interpreted the data. J.F.V.-C., L.M. and B.B. provided key resources. T.Q., M.M. and B.B. prepared the original manuscript. All authors reviewed and edited the manuscript. T.Q. and L.A. prepared the figures. B.B, L.M., and B.A.T. supervised the study. L.M. and B.B. administered the project. B.A.T., L.M. and B.B. acquired funding.

## Code availability

All scripts used for downstream analysis and to reproduce all figures are in https://github.com/BEBlab/SOD1-ddPCA and https://github.com/lashcroftuow/FLIM-analysis-code.

## METHODS

### Protein residues nomenclature

Given this work is focused at the protein level, annotations of SOD1 residues do not take into account the N-terminus Methionine. Thus, the starting residue here is referred to Alanine 1.

### Library design

The SOD1 coding sequence was split into 3 equal-length fragments (159 bp each). Libraries were designed for each fragment, resulting in 3 libraries of 200 nt each, including flanking nucleotide sequences as constant regions (19-20nt length depending on the library). Each fragment was designed to have a 3 amino acid overlap with the adjacent fragment for downstream fitness scores normalization (see Abundance/binding scores and error estimates).

Each SOD1 library contains 2000 mutations, including 1007 single amino acid substitutions, 30 random synonymous, 1 WT, and for each library, respectively, 16, 17 and 14 non-sense (every 4 AA), 894, 893 and 899 random single amino acid insertions, and 52, 51 and 48 single amino acid deletions. Non-sense mutations were forced to always be included in the overlapping region between libraries. Single amino acid and synonymous substitutions were prioritized to occur by more than 2 nt changes, and when not possible, a second random synonymous mutation in another position of the same sequence was added. All library sequences were codon optimized using the *Saccharomyces cerevisiae* codon usage. The 3 SOD1 libraries were synthesized by Twist bioscience as oligopools.

### Cloning plasmid preparation

In order to fuse each library with the rest of the WT sequence, “empty vectors” carrying the SOD1 WT sequence except the respective library were created. SOD1 chunks were amplified by a PCR of 25 cycles (Q5 high-fidelity DNA polymerase, NEB) with primers annealing to their respective part of the chunk (Primers 1-8, Supplementary Dataset S1). These primers also included constant regions for downstream fusion of SOD1 chunks with the plasmid and the respective library. In parallel, a cloning vector of ∼2 kb was linearized by a PCR of 30 cycles (Primers 9-14, Supplementary Dataset S1) (Q5 high-fidelity DNA polymerase, NEB) and 1ul of DpnI was added to each 50 ul reaction for 10h (Fast Digest DpnI). SOD1 chunks were purified by column (MinElute PCR Purification Kit, Qiagen), and the linearized nicking vector was purified from a 1% agarose gel (QIAquick Gel extraction Kit, Qiagen). The SOD1 chunks were ligated into 300 ng of the linearized nicking vector in a 1:10 (vector:insert) ratio by a Gibson approach with 3h of incubation at 50°C. Each gibson product was transformed into Max efficiency DH5-α competent *E.coli* (NEB) by heatshock. Cells were recovered in 450 ul SOC medium for 1h and plated overnight in LB spectinomycin plates. Bacteria colonies were incubated overnight in LB spectinomycin liquid media and 3 ml of cultures were harvested to purify the empty vectors with a miniprep (QIAprep Miniprep Kit, Qiagen). Correct assembly of SOD1 chunks and vector was confirmed by Sanger sequencing. Finally, each empty vector was linearized at their respective sites for downstream cloning of the SOD1 libraries (Primers 9-14, Supplementary Dataset S1), and DpnI was added.

### Abundance and binding PCA plasmids preparation

In the aPCA pGJJ162 plasmid the DHFR3 fragment is fused to the C-terminus of the target protein and the expression is driven by the CYC promoter, while the DHFR1,2 fragment is expressed at high levels as is driven by the GPD promoter. In the bPCA pGJJ159 plasmid, both DHFR3 and DHFR1,2 fragments are fused to the C-terminus of the target proteins, both fusions are expressed at the same levels, driven by the CYC promoter.

pGJJ162 was digested with the NheI and HindIII restriction enzymes for 3 h at 37°C, followed by an enzyme inactivation for 10 min at 80°C. pGJJ159 was first digested with the SpeI and BamHI enzymes and ligated with a re-coded SOD1 WT sequence (oligonucleotide 21 and primers 22 and 23, Supplementary Dataset S1), and then digested with the NheI and HindIII. After each digestion vectors were dephosphorylated for 1h at 37°C with 1 ul of FastAP enzyme.

### Library cloning

10 ng of each library pool were amplified by a PCR of 12 cycles (Q5 high-fidelity DNA polymerase, NEB) with primers annealing to the constant regions (Primers 15-20, Supplementary Dataset S1). The products were incubated with 2 ul of ExoSAP (Affymetrix) for 1 h at 37°C and purified by column (MinElute PCR Purification Kit, Qiagen). SOD1 libraries were ligated into 300 ng of their respective linearized empty vectors in a 1:10 (vector:insert) ratio by a Gibson approach with 3h of incubation at 50°C. Each Gibson reaction was dialyzed for 3 h on a membrane filter (MF-Millipore 0.025 μm membrane, Merck), followed by a vacuum concentration. The product was transformed into 10-beta Electrocompetent *E. coli* (NEB), by electroporation at 2.0 kV, 200 Ω, 25 μF (BioRad GenePulser machine). Cells were recovered in SOC medium for 1 h and grown overnight in 50 ml of LB spectinomycin medium. A small number of cells were also plated in LB spectinomycin plates to assess transformation efficiency. A total of >200,000 transformants were estimated, meaning that each variant in each library is represented >100 times. 5 ml of overnight cultures were harvested to purify the SOD1 libraries with a mini prep (QIAprep Miniprep Kit, Qiagen). Each library miniprep was digested with the NheI and HindIII enzymes for 3h at 37°C, followed by enzyme inactivation for 10min at 80°C, purified from a 1% agarose gel (QIAquick Gel extraction Kit, Qiagen) and ligated with their respective PCA plasmids in a 1:5 (plasmid:insert) ratio with the T4 DNA ligase enzyme for 10h. Each ligation was dialyzed for 2 h on a membrane filter (MF-Millipore 0.025 μm membrane, Merck) and vacuum concentration. The product was transformed into 10-beta Electrocompetent *E. coli* (NEB), by electroporation at 2.0 kV, 200 Ω, 25 μF (BioRad GenePulser machine). Cells were recovered in SOC medium for 1 h and grown overnight in 50 ml of LB ampicillin medium. A small number of cells were also plated in LB ampicillin plates to assess transformation efficiency. A total of >200,000 transformants were estimated, meaning that each variant in each library is represented >100 times. 5 ml of overnight cultures were harvested to purify the SOD1 libraries with a mini prep (QIAprep Miniprep Kit, Qiagen).

### Yeast transformation

*Saccharomyces cerevisiae* BY4742 (MATα his3Δ1 leu2Δ0 lys2Δ0 ura3Δ0) cells were used in all yeast experiments in this study. Yeast cells were transformed with the SOD1 libraries in three biological replicates. Per replicate, an individual colony was grown overnight in 3 ml YPDA medium at 30 °C and 4 g. Cells were diluted in 50 ml to OD600 = 0.2 and grown for 4–5 h. When cells reached the exponential phase (OD∼0.8–0.9) they were harvested at 3000 × g for 5 min, washed with 50 ml milliQ, centrifuged at 3000 × g for 5 min and washed with 25 ml SORB buffer (100 mM LiOAc, 10 mM Tris pH 8.0, 1 mM EDTA, 1 M sorbitol). Cells were resuspended in 1.4 ml of SORB and incubated 30 min on an orbital shaker. After incubation, 1000 ng of library and 30 ul of ssDNA (UltraPure, Thermo Scientific) were added and incubated for 5 min at room temperature and then 10 min on an orbital shaker. 6 ml of YTB-PEG (100 mM LiOAc, 10 mM Tris pH 8.0, 1 mM EDTA, 40% PEG 3350) and 600 ul of DMSO were added to the cells. Heat-shock was performed at 42°C for 20 min in a liquid bath with intermittent shaking. Cells were harvested and incubated in 50 ml of recovery medium (YPDA medium + Sorbitol 0.5 M) for 1 h at 30°C. Cells were harvested and grown in 50 ml plasmid selection medium (-URA, 2% glucose) for 50 h at 30 °C. A small amount of cells were also plated in plasmid selection solid medium to assess transformation efficiency. 200,000-400,000 transformants were estimated for each biological replicate, meaning that each variant in the library is represented at least 100 times. After 50 h, cells were diluted in 50 ml plasmid selection medium to OD= 0.05 and grown exponentially for 15 h. Finally, the culture was harvested and stored at −80 °C in 25 % glycerol.

### Selection assays

*In vivo* selection assays were performed in three independent biological replicates for each SOD1 library. For each replicate, cells were diluted from the 50 h culture at OD = 0.05 and grown until exponential for 12 h. When cells reached exponential, they were diluted in 50 ml plasmid selection media (-URA -ADE -MET, 2% glucose) at OD = 0.2. After 6 h, cells were diluted in 50 ml plasmid selection media (-URA -ADE -MET +MTX, 2% glucose) at OD = 0.04. The 40ml that were not used for dilution were collected as input pellets. Selection in MTX was performed until cells reached exponential (∼22h) and the 50 ml were collected as output pellets. Both input and output pellets were stored at −20 °C before DNA extraction. Three input and three output samples were processed for sequencing.

### DNA extraction and sequencing library preparation

Input and output pellets were resuspended in 0.5ml extraction buffer (2% Triton-X, 1% SDS, 100 mM NaCl, 10 mM Tris-HCl pH 8, 1 mM EDTA pH 8). They were then frozen for 10 min in an ethanol-dry ice bath and heated for 10 min at 62 °C. This cycle was repeated twice. 0.5ml of phenol:chloroform:isoamyl (25:24:1 mixture, Thermo Scientific) was added together with glass beads (Sigma). Samples were vortexed for 10 min and centrifuged for 30 min at 2000 × g. The aqueous phase was then transferred to a new tube, and mixed again with phenol:chloroform:isoamyl, vortexed and centrifuged for 45 min at 2000 × g. Next, the aqueous phase was transferred to another tube with 1:10 V 3 M NaOAc and 2.2 V cold ethanol 96% for DNA precipitation. After 1 h at −20 °C, samples were centrifuged and pellets were dried overnight. The following day, pellets were resuspended in 0.3 ml TE 1X buffer and treated with 10 μl RNAse A (Thermo Scientific) for 30 min at 37 °C. DNA was finally purified using 10 μl of silica beads (QIAEX II Gel Extraction Kit, Qiagen) and eluted in 30 μl elution buffer. Plasmid concentrations were measured by quantitative PCR with SYBR green (A25742, Applied biosystems) and primers annealing to the origin of replication site of the pGJJ162 and PGJJ159 plasmids at 58 °C for 40 cycles (Primers 139 and 140, Supplementary Dataset S1). The library for high-throughput sequencing was prepared in a two-step PCR (Q5 high-fidelity DNA polymerase, NEB). In PCR1, 50 million of molecules (SOD1 libraries) were amplified for 15 cycles with frame-shifted primers with homology to Illumina sequencing primers (Primers 24-66, Supplementary Dataset S1). The products were treated with ExoSAP treatment (Affymetrix) and purified by column purification (MinElute PCR Purification Kit, Qiagen). They were then amplified for 12 cycles in PCR2 with Illumina-indexed primers (Primers 66-138, Supplementary Dataset S1). For each of the SOD1 libraries, the six samples (one input-output pair per biological replicate) were pooled together equimolarly and the final product was purified from a 2% agarose gel with 20 μl silica beads (QIAEX II Gel Extraction Kit, Qiagen). The libraries were sent for 150 bp paired-end sequencing in an Illumina NextSeq500 sequencer at the CRG Genomics core facility. In total, >10 million paired-end reads were obtained for each of SOD1 libraries, representing >1000x read per variant coverage.

### Individual variant testing

SOD1 single mutants were obtained by PCR mutagenesis of the SOD1 WT with 25 cycles (Q5 high-fidelity DNA polymerase, NEB) (Primers 141-170, Supplementary Dataset S1). The products were purified and cloned in bacteria in the same way as the SOD1 libraries. Correct mutagenesis was confirmed by Sanger sequencing and plasmids were used to transform yeast BY4742. Yeast colonies were grown overnight in 3ml of Gluc -URA and diluted to OD 0.3 in Gluc -URA -ADE - MET the morning after. Selection with MTX started at OD 0.1 until saturation. The growth rate of each variant was measured in triplicates.

### Data processing

FastQ files from paired end sequencing of the libraries were processed using DiMSum^57^, an R pipeline for analyzing deep mutational scanning data. 5′ and 3′ constant regions were trimmed, allowing a maximum of 20% of mismatches relative to the reference sequence. Sequences with a Phred base quality score below 30 were discarded. Non-designed variants were also discarded for further analysis, as well as variants with input reads below the threshold suggested by DimSum in all of the replicates (the input counts threshold for libraries were 100 for aPCA and 10 for bPCA).

### Abundance/heterodimerization scores and error estimates

The DiMSum package (https://github.com/lehner-lab/DiMSum)^57^ was also used to calculate both abundance and heterodimerization scores and their error estimates for each variant in each biological replicate. For each variant (*i*) in replicate (*r*), an enrichment score (ES) was first computed as the natural logarithm of the ratio between its normalized sequencing read counts in the output (collected after selection) and input (collected before selection) samples:

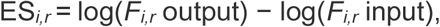

where *F* represents the sequencing read frequency of a given variant in each library. Likewise, the enrichment score for the SOD1 WT sequence was calculated as:

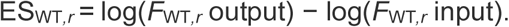

The abundance/heterodimerization score for each variant was then defined as the difference between its enrichment score and that of the WT in the same replicate:

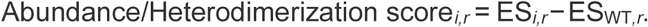

Scores for each variant were merged across biological replicates using error-weighted mean and centered to the WT scores. All scores and associated error estimates are available in Supplementary Dataset S2.

All the mutation types included in the SOD1 library (substitutions, insertions and deletions) were used to center, and categorize the abundance and heterodimerization data.

Each dataset was relatively normalized to the median score of stop mutations. Sigma values were normalized to the interquartile range (relative to each assay) and variants with a normalized sigma above a cut-off of 0.3 were considered as noisy variants and excluded from further analysis.

### Abundance/heterodimerization scores classification

Abundance and heterodimerization categories were defined by following both biological and statistical criteria. First, a threshold for synonymous mutations was calculated and defined as the synonymous score that separates the 95% of mutations with the highest abundance/heterodimerization from the 5% of mutations with the lowest abundance/heterodimerization (−0.19 and -0.05, respectively). The same reasoning was applied to calculate a threshold for stop mutations in both abundance and heterodimerization (−0.87, - 0.86, respectively). Second, a False Discovery Rate (FDR) was tested against both synonymous and stop thresholds. Combining the two criteria, mutations were classified as “wt-like” when the score was higher than the synonymous threshold and their sigma non-significantly different from synonymous; “low” when the score was lower than the synonymous but higher than the stop threshold, and the sigma significantly different from both synonymous and stops; “stop-like”, lower than the stop threshold and not different from stops; “High abundance/heterodimerization” mutations were defined as those with a score higher than the median of synonymous mutations.

### Statistics and reproducibility

Based on transformation efficiency, each variant in the designed libraries was expected to be represented at least 100x, a coverage that was maintained at each step of the library cloning, yeast transformation, selection experiments and library preparation for sequencing. In terms of sequencing, reads that did not pass the QC filters using the DiMSum package were excluded (https://github.com/lehner-lab/DiMSum)^57^. The experiments were not randomized. The investigators were not blinded to allocation during experiments and outcome assessment

### Data modelling with LOWESS

Data was modeled using all variants in the SOD1 dataset, including substitutions, insertions, deletions and premature stops. The loess function in R was used, adjusting the weights of WT-like and non-sense scores to 1e04 each, and span = 0.5. Heterodimerization residuals were calculated as the difference between the observed heterodimerization score and the LOWESS-predicted heterodimerization score given the abundance score. Span values, ranging from 0.1 to 1, were evaluated. A span = 0.5 provided the minimal bias of residuals as a function of the fitted scores.

### Definition of residual categories

Mutations with significant residual were considered as those whose residual was higher than 2*MAD (Median Absolute Deviation). Mutations with significant residual were split into two categories, according to the residual sign: high positive and high negative residual.

### Criteria for defining allosteric positions/mutations

Allosteric positions were defined by employing a similar approach reported in^58,59^. The median HD residual by position and the minimal distance to the dimer interface were used to define allosteric positions. A first threshold was defined as the median of HD residuals of all the dimer interface residues (from substitutions and indels) and a second threshold as the highest distance of the dimer interface residues (A4). Allosteric positions were defined as those where their median HD residual was higher than the median HD residuals of dimer interface residues and not being part of the dimer interface (distance to dimer interface>highest distance to dimer interface). The same criteria was used to define allosteric mutations.

### Variant effect predictors

The VEPs scores were retrieved from https://alphamissense.hegelab.org/ ^60^ (Alphamissense), https://huggingface.co/spaces/ntranoslab/esm_variants^61^ (ESM-b1), https://sites.google.com/site/revelgenomics ^62^ (REVEL), and https://evemodel.org/ ^63^ (EVE).

### 3D protein visualization

3D SOD1 structural analysis and visualization were performed with ChimeraX^64^. Abundance, heterodimerization and heterodimerization residual scores were transformed to b-factors and used to map mutational impacts in SOD1 (PDB: 2V0A^65^). Contacting residues were identified with the “Contacts” function, with distances between the center of the pair of atoms being lower than 5Å.

### Relative solvent accessible surface area (rSASA) predictions

rSASA for each SOD1 (PDB:2V0A) position was calculated for both monomeric (monomer rSASA) and dimeric (dimer rSASA) states with ChimeraX^64^. Positions with a rSASA (whether in the monomer or dimer) lower than 25 were considered as buried residues, while positions with a rSASA > 25 were considered as surface-exposed residues.

### Definition of SOD1 pockets

The CASTpFold^66^ server was used to predict pockets in SOD1 with the “run online” option, and using the PDB 2V0A as an input.

### Plasmid preparation for mammalian cell culture experiments

Plasmids used for FLIM experiments consist of each SOD1 variant with N-terminal AcGFP and C-terminal mCherry, cloned into a pcDNA3.1(+) backbone and were synthesized and cloned by Genscript (USA). Plasmids used for analysis of SOD1 inclusions, intensity and impact on cell survival are composed of each of the SOD1 variants with C-terminal EGFP, cloned into pEGFP-N1 by Genscript (USA). All plasmids were heat transformed into chemically competent *E. coli* DH5α cells and purified from bacterial culture using Genejet miniprep kits (ThermoFisher Scientific) as per manufacturer’s instructions.

### Mammalian cell culture

HEK293T cells were maintained in high glucose Dulbecco’s modified eagle medium (DMEM) with 10% (*v/v*) heat inactivated fetal bovine serum and 1× GlutaMAX. Cells were plated out at a confluency of 20% prior to experimentation. NSC-34 cells were maintained in DMEM/F12 with 10% (*v/v*) heat inactivated fetal bovine serum and 1× GlutaMAX and plated at a confluency of 40%. For fluorescence lifetime imaging (FLIM) experiments HEK293T cells were transfected 24 h post-plating with 0.1 µg of plasmid DNA per chamber of an 18-well µ-slide (Ibidi), using Lipofectamine 3000, according to the manufacturer’s instructions. Meanwhile for aggregation, cell counting, and intensity measurement experiments, HEK293T and NSC-34 were plated into 24-well plates and transfected with 0.5 µg of plasmid using Lipofectamine 3000.

### Live cell fluorescence lifetime imaging microscopy

Cells were imaged live (37 °C, 5% CO_2_) using a SP8 Fast Lifetime CONtrast (FaLCon) confocal fluorescence microscope (Leica Microsystems) 48 h post-transfection. AcGFP and mCherry were excited by a pulsed (40 MHz) white light laser (85% power), respectively, and emission was detected at 498–551 nm (AcGFP) and 620–784 nm (mCh) using a 10× air objective. Images were obtained with a 16-bit depth, 256 × 256 pixel resolution, 200 Hz scan speed in bidirectional mode, a pinhole of 2 airy units and 5× digital zoom. Laser transmission was altered between images to maintain adequate signal and reduce over-exposure of pixels (minimum photon count/frame > 7,500 photons). As such, the laser power varied between 0.04–0.8 % (Supplementary Dataset S3) due to differential variant abundance. For each field of view, 30 frame repetitions were acquired. AcGFP fluorescence lifetime images were represented as a colour spectrum from blue (1.4 ns) to red (2.8 ns) using Las X FLIM/FCS software (Leica Microsystems). Additionally, the ‘fast FLIM’ function within the software was used to collate and export the density of AcGFP fluorescence lifetimes from 0–25 ns for custom analysis in Python.

### Analysis of FLIM data

For all expressed FLIM plasmids, the AcGFP fluorescence lifetime frequency distribution histograms were truncated to disregard lifetimes outside of a relevant range for AcGFP signal, which were therefore only reporting noise/background. This was automated for consistency between replicates by filtering out the lifetimes for photon densities which were less than 0.5% of the maximum AcGFP signal in that treatment. The histograms were normalized such that the area under the curve was equal to 1, and converted to a cumulative frequency histogram. The cumulative histogram was then fitted to a Boltzmann-sigmoid function using the Python package scipy ‘curve_fit’ function, and parameters were extracted. The V50 describes the median lifetime for each replicate and the mean ± SEM V50 was determined for each group.

### SOD1 microscopy

Cells were imaged using a DMi8 fluorescence microscope (Leica Microsystems) 24, 48, and 72 h post-transfection. EGFP was excited by an arc mercury lamp, and emission was detected using a GFP filter cube using a 10× air objective. Images were obtained with a 16-bit depth, 2048 × 2048 pixel resolution. Images were then analyzed as previously described (McAlary *et al.*, 2022; Shephard *et al.*, 2024) to extract information about inclusion formation, cell number, and SOD1 levels. Images were analyzed using Cellpose3^67^ to first identify all transfected cells, then CellProfiler (4.2.8) was used to measure EGFP intensity, texture, radial intensity and granularity^68^. These metrics were used to train a CellProfiler Analyst^69^ model to identify the percentage of cells with inclusions. Finally, transfection efficiency at 24 and 72 h was determined by comparing the proportion of total cells that were fluorescent. The relative survival rate of cells containing each variant was calculated via comparison of the transfection efficiency at 24 and 72 h normalized to SOD1^WT^.

### Clinical phenotypes data collection

Variants with clinical phenotypes annotations were collected from the literature (see Supplementary Dataset S8). Given the diversity in the clinical annotations, each phenotype was grouped in the following way:

<u>Penetrance</u>: annotations with “complete” or “high” penetrance were defined as “Complete + high penetrance”. Annotations with “incomplete” or “greatly reduced” were defined as “Incomplete penetrance”.

<u>Survival time</u>: annotations with “long”, “longest (44-45 y)”, “>10 years” and “11 years” were defined as “Long survival time”. Annotations with “short”, “shortest (2-3 m)” and “<3 years” were defined as “Short survival time”.

<u>Progression</u>: the annotations from the literature (“fast” and “slow”) were used.

### Clinical calibration of abundance and heterodimerization scores

A validation set of well-characterized pathogenic, likely pathogenic, benign, and likely benign *SOD1* variants were collected after classification using adapted American College of Medical Genetics and Genomics/Association of Molecular Pathology (ACMG/AMP) variant classification criteria^70^ without the contribution of functional assay evidence. Variants were collected from public genetic databases (ClinVar^37^, ALSoD https://alsod.ac.uk/, LOVD https://www.lovd.nl/) in July 2024. Using a previously adapted Bayesian approach^54^ the likelihood ratios (LR) for normal and abnormal function were calculated based on the categorization of the abundance and dimerization scores (WT-like, low, stop-like). The LRs for normal and abnormal function were converted to a BS3 and PS3 strength, respectively using the previously published cut-offs^71^.

### Clinical reference sets for comparison with Variant Effect Predictors

A reference set containing synonymous variants was used to test abundance and heterodimerization scores at classifying deleteriousness. However, given that VEPs do not provide estimates for synonymous variants, we employed the reference set used by Axakova et al.^35^ to compare the performance of abundance and heterodimerization with Alphamissese, EVE, REVEL, and the VAMP-seq data at classifying deleteriousness. This reference set is composed of 84 pathogenic variants and 100 “proxy” benign variants (i.e., variants available in gnomAD with no clinical annotations at the time of publication).

---

**Supplementary figure 1.**
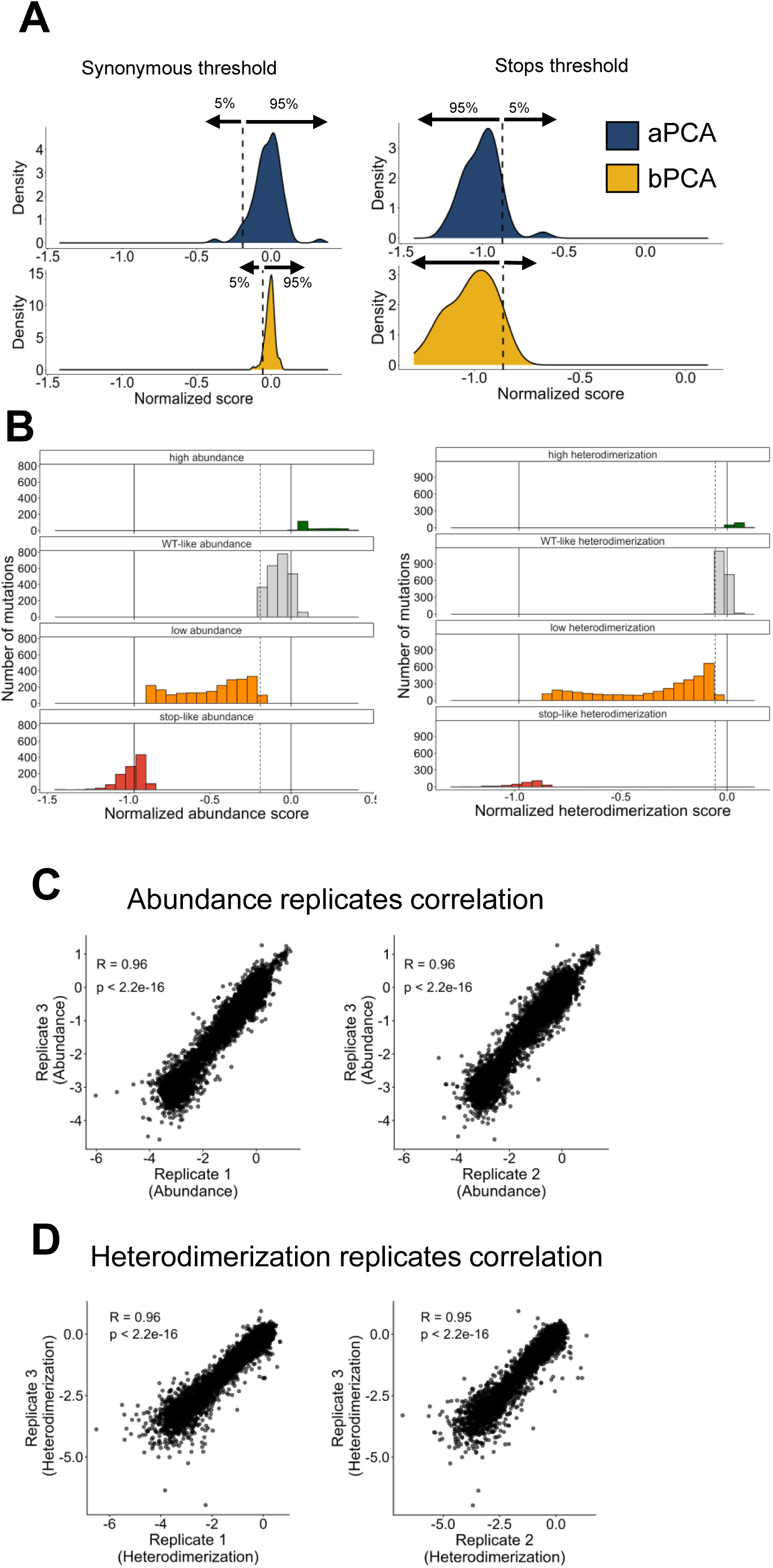
Abundance and heterodimerization data categorization. **(A)** Synonymous (left) and premature stops (right) score distributions in the aPCA (blue) and bPCA (yellow). Dashed lines indicate the 95th percentile. **(B)** Scores distribution of FDR categories in abundance (left) and heterodimerization (right). Solid lines indicate the premature stops modes (score ∼-1) and WT score (score=0); dashed lines indicate synonymous thresholds (95th percentile). Scatter plot of abundance replicates **(C)** and heterodimerization replicates **(D).** Pearson’s coefficients and p-values are indicated.

**Supplementary figure 2.**
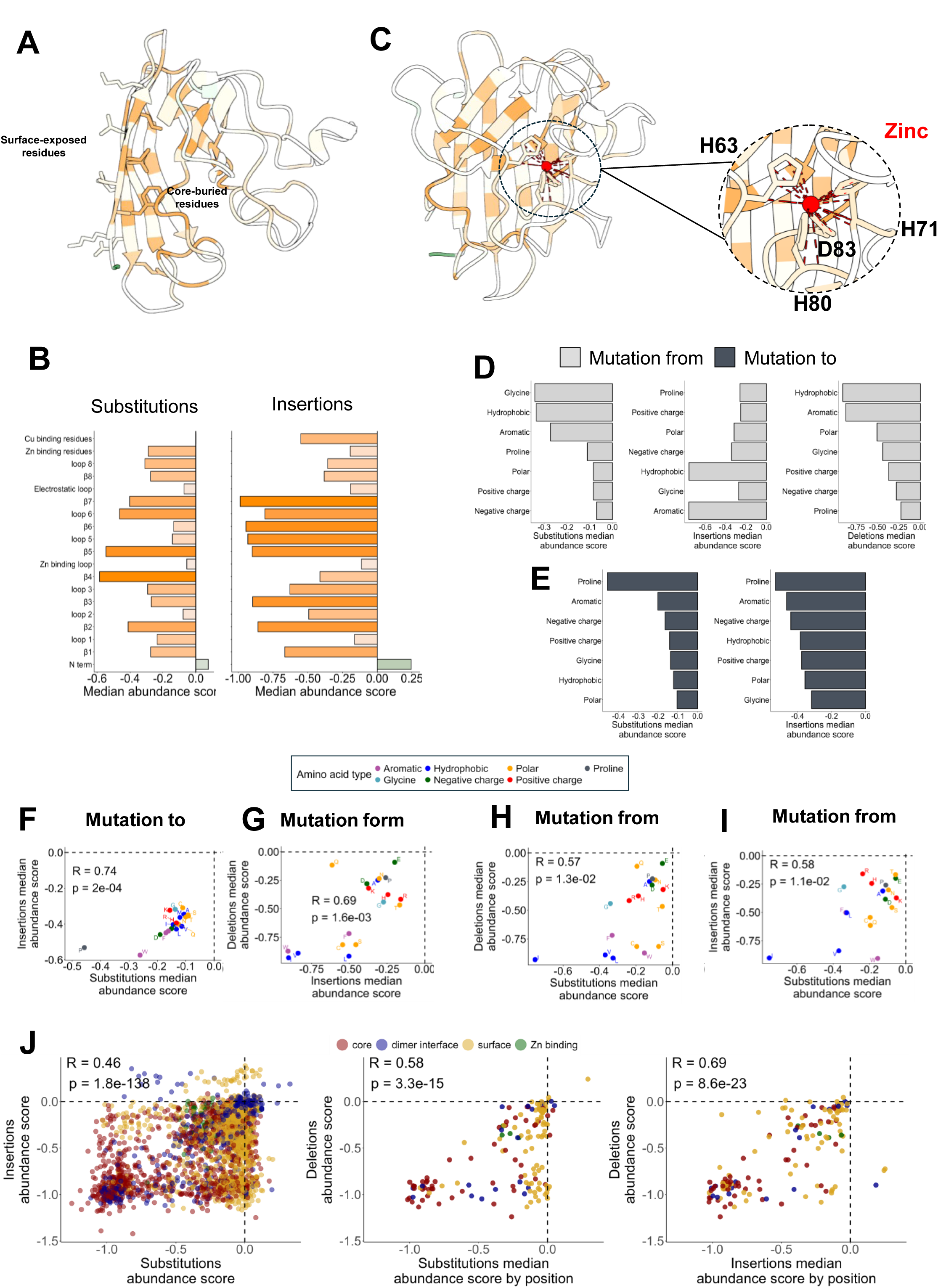
Extended analysis of the SOD1 abundance data. **(A)** SOD1 3D monomer structures (PDB: 2V0A) colored by the median abundance scores per positions of substitutions, with an example of surface-exposed and core-buried residues represented as sticks. **(B)** Bar plots showing the median mutational abundance score of SOD1 secondary structures and functional residues for substitutions (left) and insertions (right). **(C)** SOD1 3D monomer structures (PDB: 2V0A) colored by the median abundance scores per substitutions positions highlighting the zinc-binding residues, represented as sticks. The zinc atom is shown as a red ball. Bar plots showing the median abundance score of substitutions (left), insertions (middle) and deletions (right) grouped by the amino acid category of the mutated residue **(D)** or the amino acid category of the mutant residue **(E)**. Scatter plots of the median abundance scores by position of substitutions and insertions **(F)**, insertions and deletions **(G)**, substitutions and deletions **(E)** and substitutions and insertions (mutated from). Amino acids are colored according to their chemical category. Pearson’s coefficients and p-values are indicated. Dashed lines indicate WT scores. **(J)** Scatter plot comparing abundance scores of substitutions and insertions (left), median abundance scores by position of substitutions and deletions (middle) and insertions and deletions (right). Dots are colored by the side-chain and functionality of residues: red = core-buried, yellow = surface-exposed, blue = dimer interface, green = zinc-binding. Pearson’s coefficients and p-values are indicated. Dashed lines indicate WT scores.

**Supplementary figure 3.**
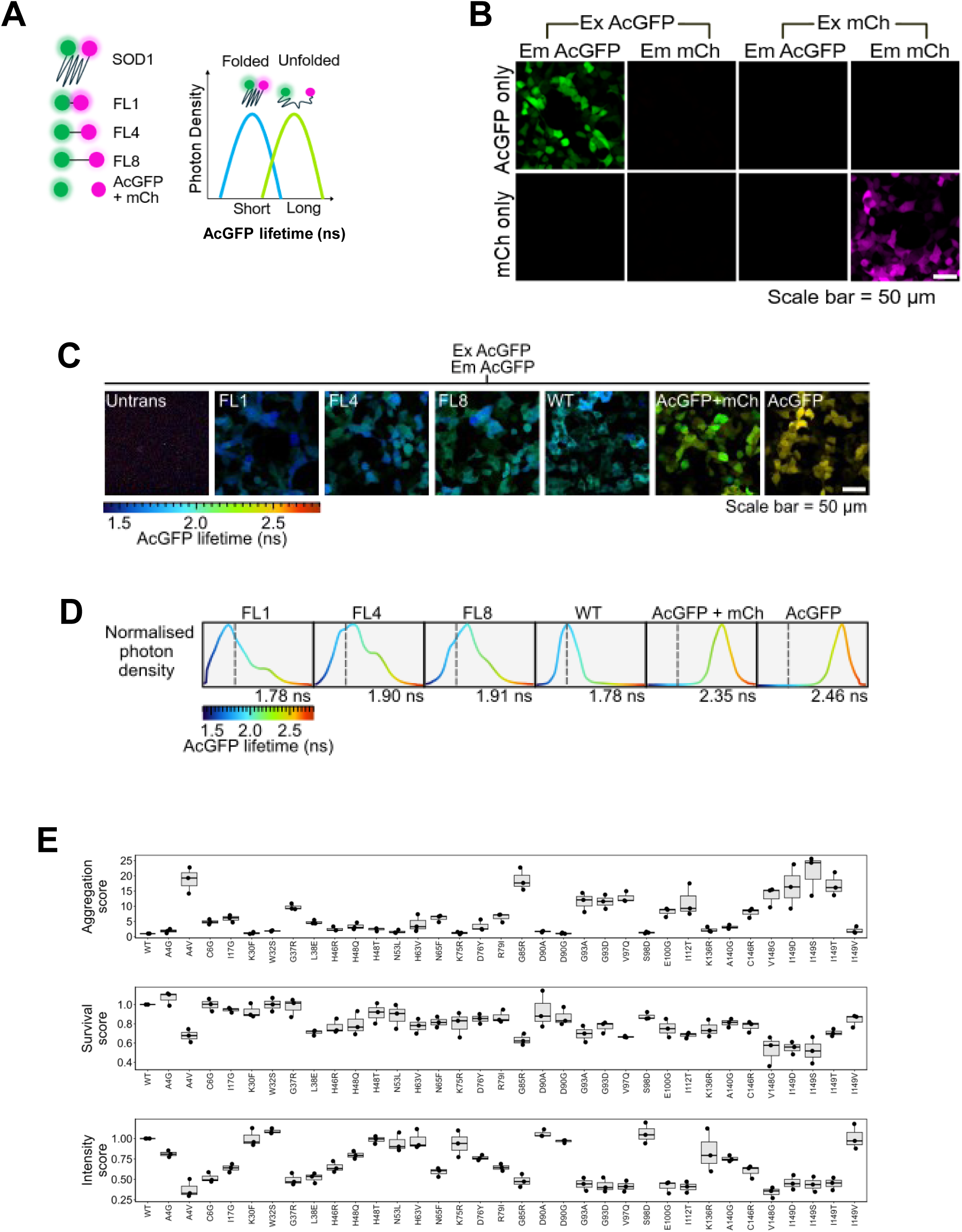
Fluorescence lifetime imaging microscopy (FLIM) is sensitive to changes in distance between AcGFP and mCherry in HEK293T cells. **(A)** Schematic representation of the AcGFP/mCh FRET-FLIM sensor used to detect conformational changes in SOD1 protein structure, in HEK293T cells. Left: AcGFP (green) is fused to the N-terminus and mCherry (magenta) is fused to the C-terminus of SOD1. Right: Expected outcome of the FRET-FILM sensor for SOD1 folding. **(B)** Spectral bleed through in each emission channel (Em AcGFP = 498–551 nm, Em mCh = 620–784 nm) following excitation at the expected excitation wavelength of each fluorophore (Ex AcGFP = 475 nm, Ex mCh = 561 nm) in cells transfected with AcGFP or mCh alone under imaging parameters specified in methods. Scale bar represents 50 µm. **(C)** Lifetime imaging of fluorescent linker constructs showing that as linker length decreases, so does the AcGFP lifetime. **(D)** Quantification of images from (C), showing lifetime distributions. Median lifetime for each treatment is indicated at the bottom right of each box and wild-type SOD1 median lifetime is shown as a grey dotted line for comparison. **(E)** Individual measurements of SOD1 variants (n=34) inclusions (aggregation), survival, and intensity. All phenotype measurements are the result of a GFP-SOD1 fusion readout. Black lines in the boxes indicate mean score and error bars indicate the standard error of the mean (SEM) (n=3)

**Supplementary figure 4.**
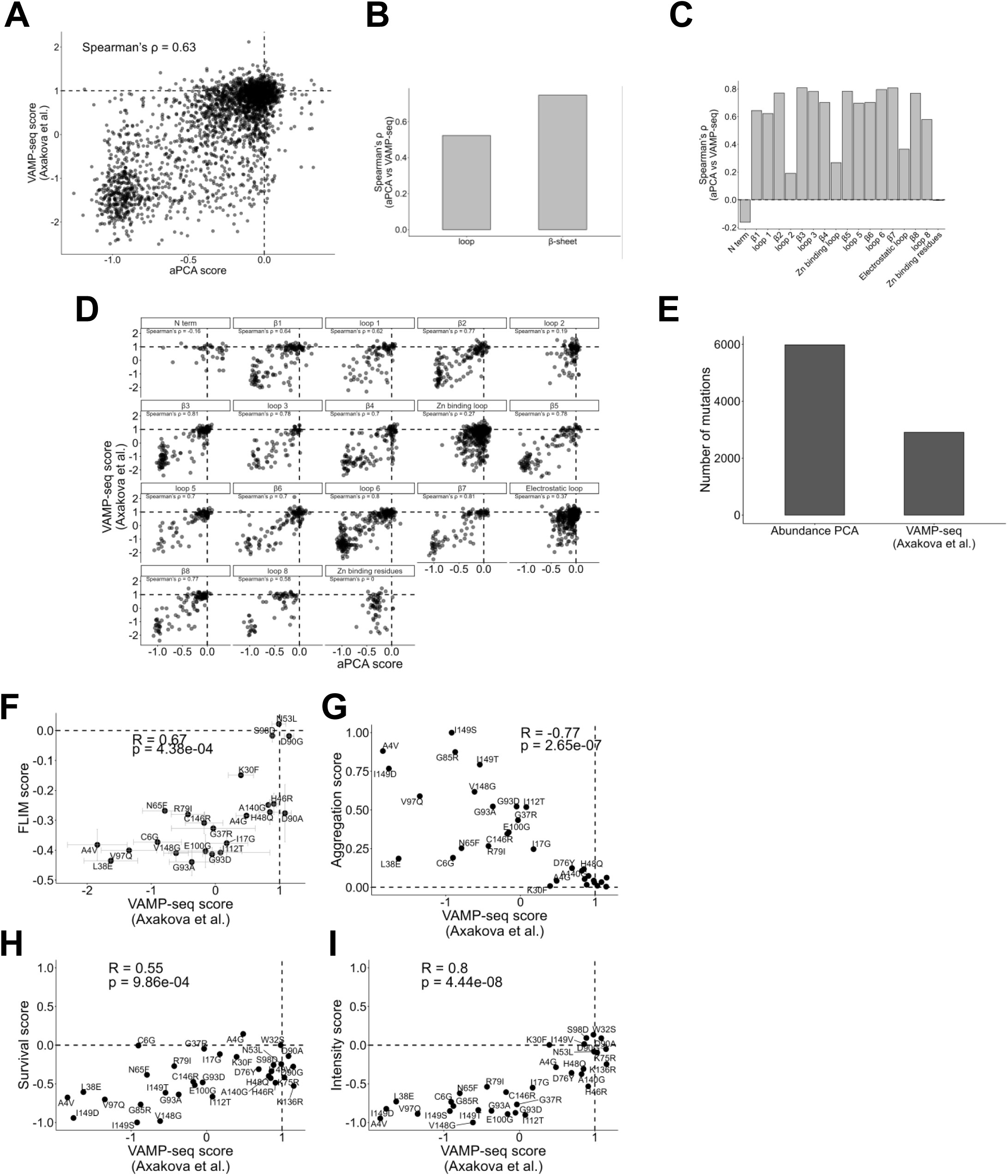
Comparison of aPCA and VAMP-seq SOD1 deep mutational scannings. **(A)** Scatter plot of aPCA (yeast) and VAMP-seq (HEK293T) scores. Dashed lines represent the WT scores of each assay (WT = 0 for aPCA, WT = 1 for VAMP-seq). Spearman’s coefficient is indicated. **(B)** Comparison of the Spearman’s coefficient of aPCA versus VAMP-seq in loops and β-strands. **(C)** Comparison of the Spearman’s coefficient of aPCA versus VAMP-seq in secondary structures. **(D)** Scatter plot matrix of the aPCA and VAMP-seq scores at each SOD1 secondary structure. Spearman’s coefficients are indicated. Dashed lines represent the relative WT scores. **(E)** Comparison of the number of mutations included in the aPCA and VAMP-seq assays. **(F-I)** Scatter plots showing the VAMP-seq scores and the validation assays scores: FLIM vs VAMP-seq **(F)**, aggregation vs VAMP-seq **(G)**, survival vs VAMP-seq **(H)** and fluorescence intensity vs VAMP-seq **(I).** Pearson’s coefficients and p-values are indicated. Dashed lines represent relative WT scores. Only the variants used for the mammalian validation assay are used for these comparisons (n=35).

**Supplementary figure 5.**
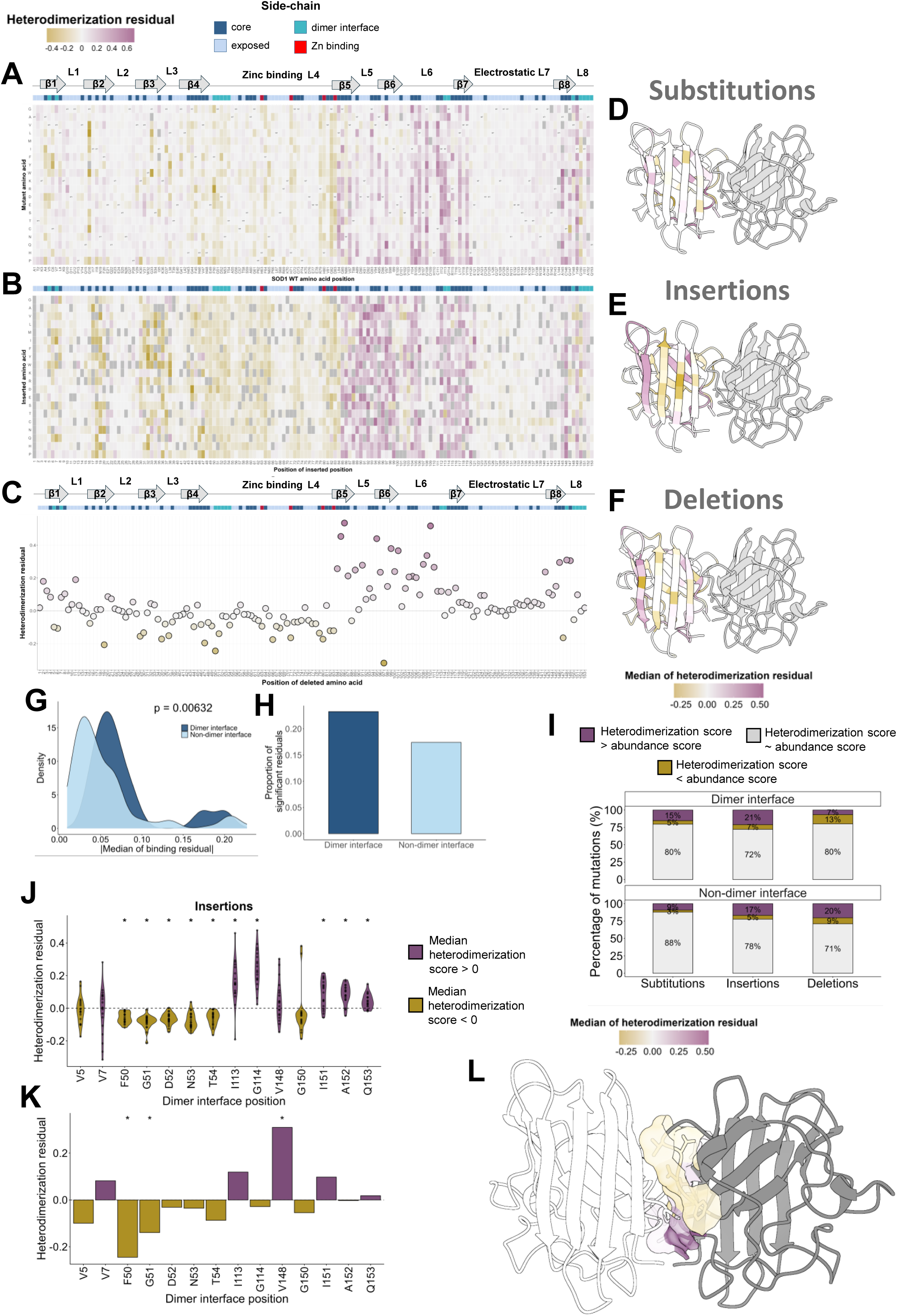
Extended analysis of the comparison between abundance and heterodimerization. Heterodimerization (HD) residual heatmap of substitutions **(A)**, insertions **(B)**, and deletions **(C).** Lineal secondary structure and residue side-chain and functionality are shown as colors at the top of each heatmap: darkblue = core-buried, lightblue = surface-exposed, cyan = dimer interface, red = zinc-binding. SOD1 3D structures (PDB:2V0A) colored by the median HD residual by position of substitutions **(D)** insertions **(E)** and deletions **(F). (G)** Comparison of the intensity of HD residual from all mutations (substitutions and indels) in dimer (darkblue) and non-dimer interface residues (lightblue). The x-axis represents the modulus of the median HD residual by position. The p-value is indicated. **(H)** Comparison of the percentage of significant HD residuals (>2MAD) in dimer and non-dimer interface residues considering all mutations. Significance is calculated with the Mann-Whitney U test. **(I)** Percentage of variants of each HD residual category for amino acid substitutions, insertions, and deletions. **(J)** Distribution of insertions HD residuals in dimer interface residues. Violins are coloured according to whether the median HD residual at a position is higher (violet) or lower (gold) than 0. Asterisks indicate positions with a significant deviation from 0 (two-sided one-sample Wilcoxon signed-rank test, Benjamini–Hochberg adjusted p-value < 0.05). The dashed line indicates heterodimerization explained by abundance (HD residual = 0). **(K)** Distribution of deletions HD residuals in dimer interface residues. Violins are coloured according to whether the median HD residual at a position is higher (violet) or lower (gold) than 0. Asterisks indicate deletions with a significant HD residual (HD residual > 2*MAD). The dashed line indicates heterodimerization explained by abundance (HD residual = 0). **(L)** SOD1 3D structure (PDB: 2V0A) where dimer interface positions are coloured by their median HD residual. The structure is flipped 180° with respect to Figure 4H.

**Supplementary figure 6.**
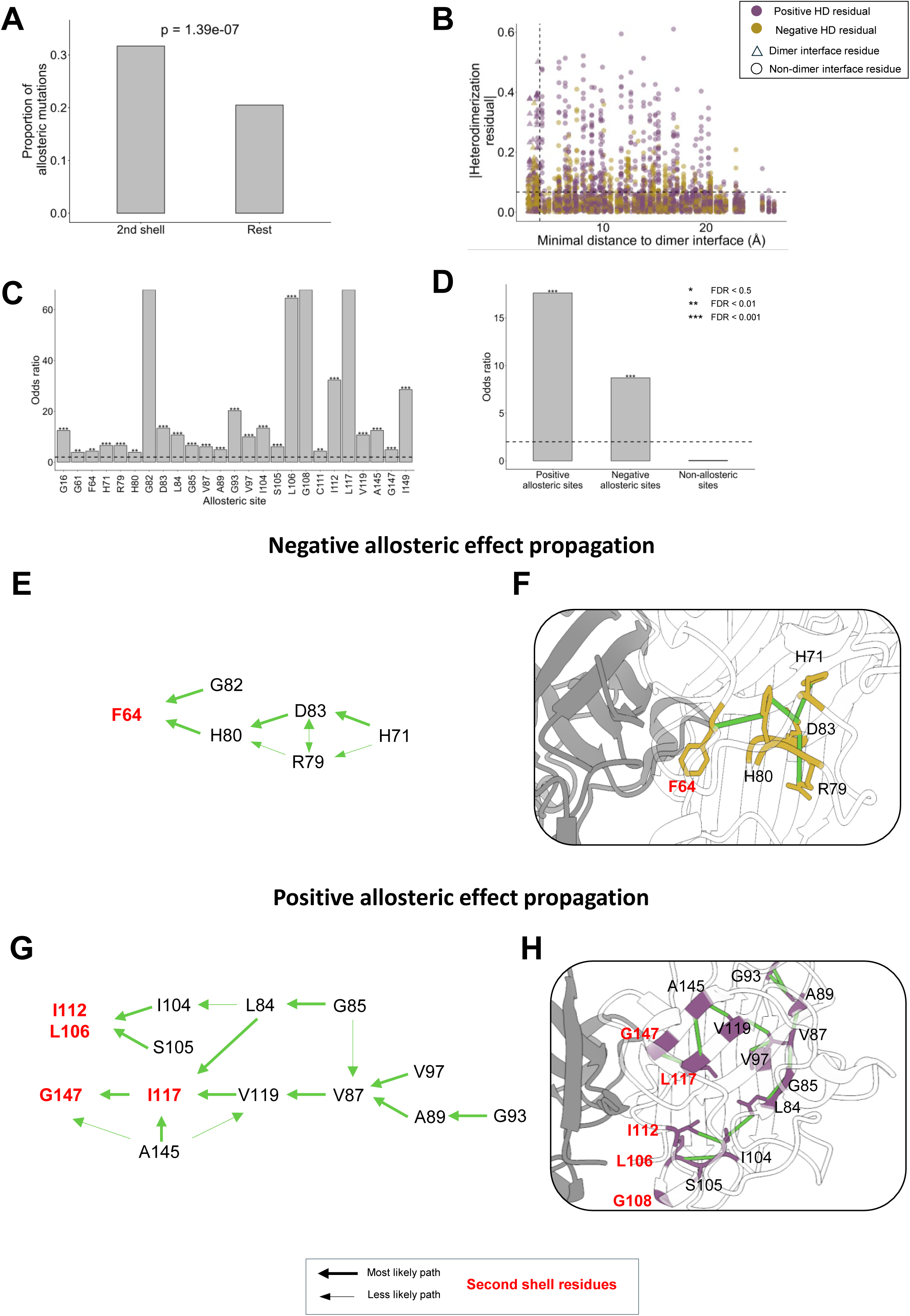
Extended analysis of the identification of SOD1 allosteric sites modulating dimerization. **(A)** Bar plot showing the proportion of allosteric mutations in second-shell residues and in the rest of residues (excluding dimer interface and second-shell residues). P-value is indicated (Chi-square test). **(B)** Relationship between per-mutation HD residual to the minimum atom distance to the dimer interface. Horizontal dashed line: absolute HD residual median of dimer interface residues; Vertical dashed line: highest minimal distance of dimer interface residues. **(C)** Enrichment of allosteric mutations at allosteric positions. Asterisks indicate statistical significance: *p<0.05; **p<0.01; ***p<0.001 (one-sided FET, Benjamini-Hochberg adjusted p-value). Dashed line indicates an odds ratio (OR)=2. **(D)** Enrichment of allosteric mutations at positive, negative and non-allosteric sites. Asterisks indicate statistical significance: *p<0.05; **p<0.01; ***p<0.001 (one-sided FET, Benjamini-Hochberg adjusted p-value).. Dashed line indicates an odds ratio (OR)=2. **(E)** Schematic representation of the hypothesized negative allosteric effect propagation. Black: allosteric sites; red: second-shell residue. The thickness of the arrows indicates the likelihood of the propagation: thick= more likely (stronger allosteric effect); thin = less likely (weaker allosteric effect). **(F)** 3D representation of the negative allosteric effect propagation, only showing residues involved in the more likely path. **(G)** Schematic representation of the hypothesized positive allosteric effect propagation. **(F)** 3D representation of the positive allosteric effect propagation, only showing residues involved in the more likely path.

**Supplementary figure 7.**
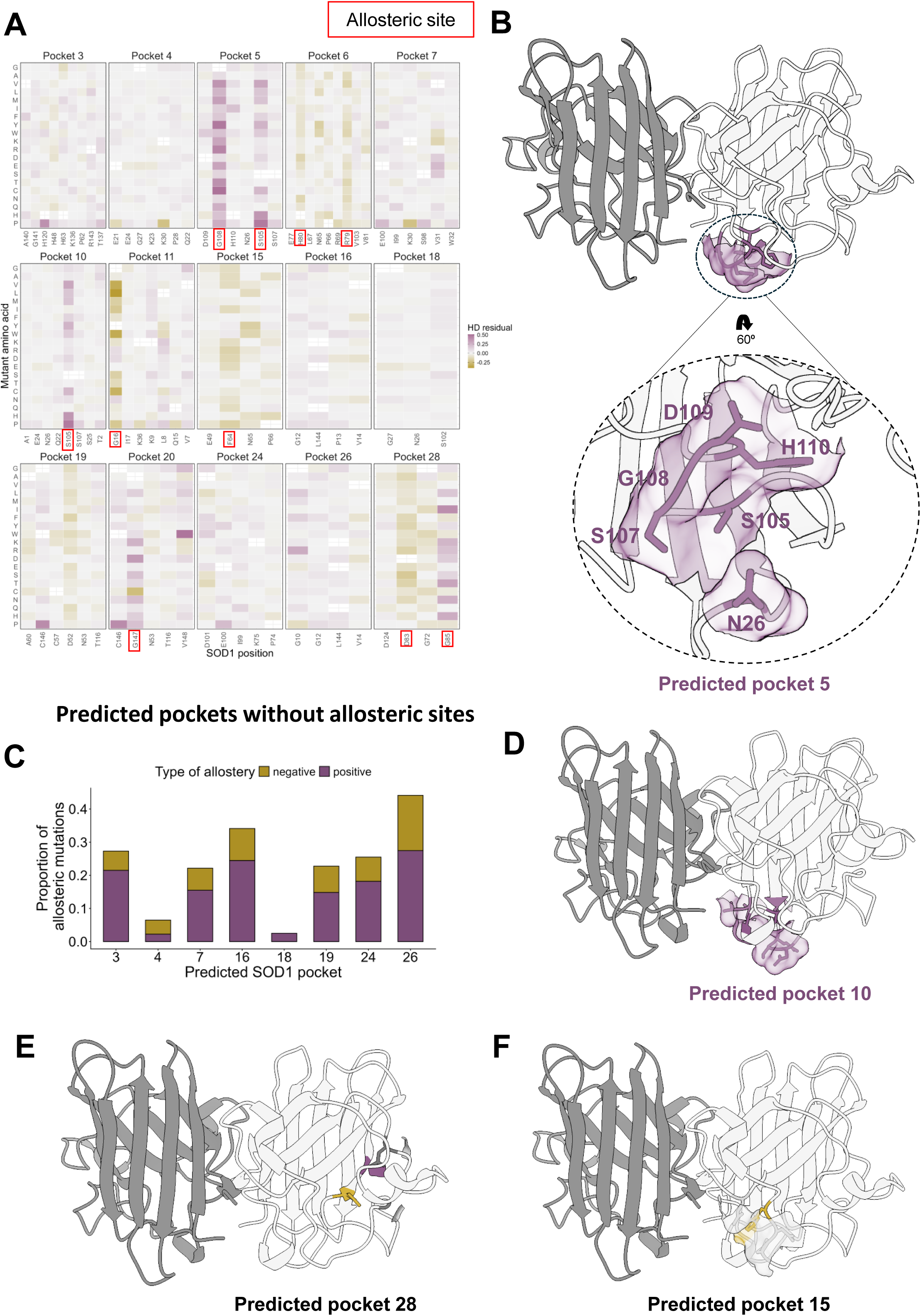
A proportion of allosteric sites cluster in predicted SOD1 pockets. **(A)** Heatmap showing the HD residuals of positions that form the predicted pockets. Allosteric sites are squared in red. **(B)** SOD1 3D structure (PDB:2V0A) highlighting the residues of pocket 5 as sticks and surface. **(C)** Proportion of allosteric mutations in predicted pockets that do not contain allosteric sites. 3D representation of the SOD1 predicted pocket 10 (enriched with positive allosteric mutations) **(D)**, and pockets 28 and 15 (containing both positive and negative allosteric mutations) **(E), (F).**

**Supplementary figure 8.**
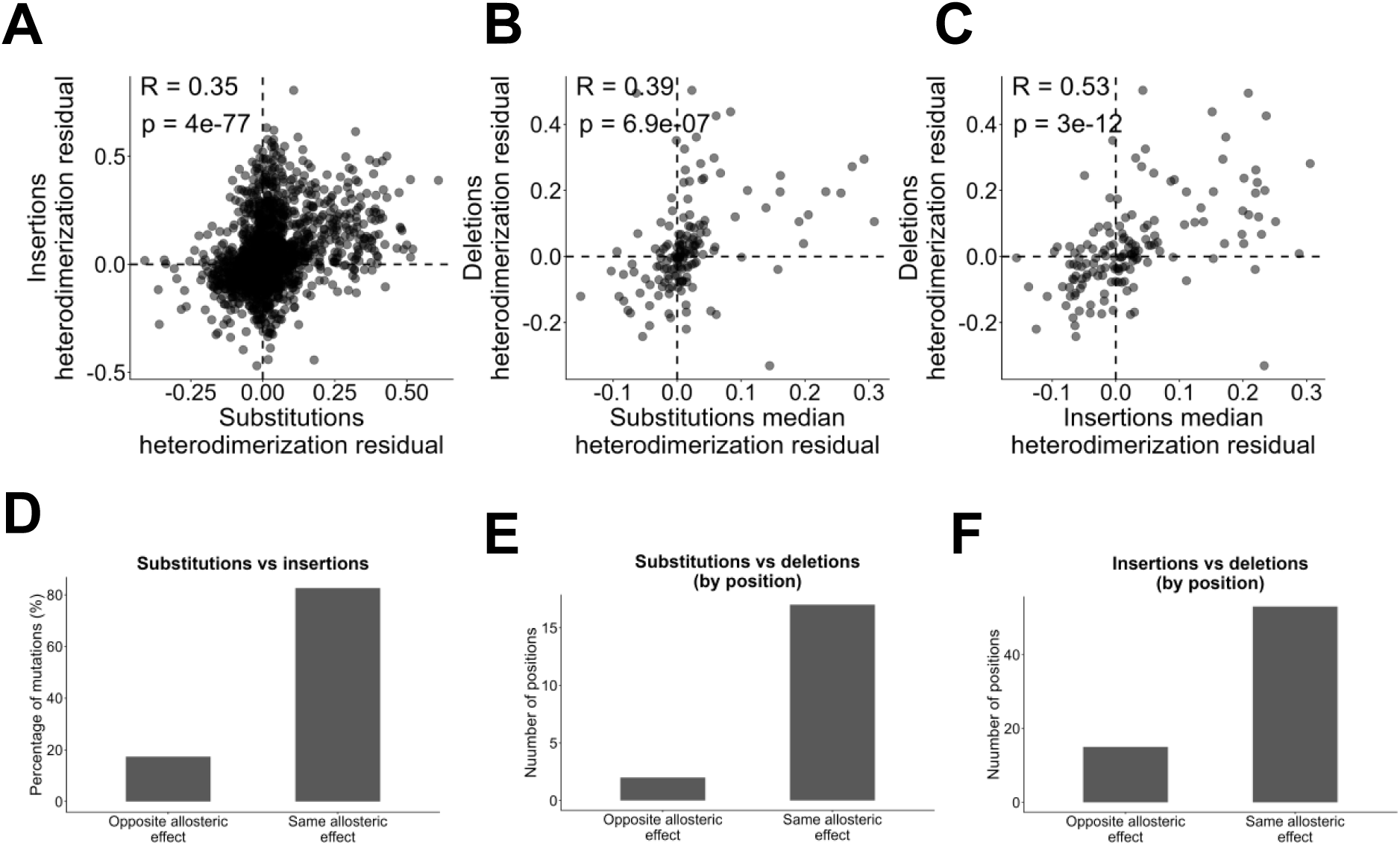
Insertions and deletions show different allosteric mechanisms with respect substitutions. Scatter plots showing the HD residuals of substitutions and insertions **(A)**, the median HD residual by position of substitutions and deletions **(B)**, and the median HD residual by position of insertions and deletions **(C)**. Pearson’s coefficients and p-values are indicated. Dashed lines at HD residual = 0 indicate the reference HD residual at which heterodimerization is explained by abundance. Percentage of mutations having the opposite allosteric effect (positive vs negative) or the same allosteric effect (positive or negative) for substitutions and insertions **(D)**, substitutions and deletions **(E)** and insertions and deletions **(F).**

**Supplementary figure 9.**
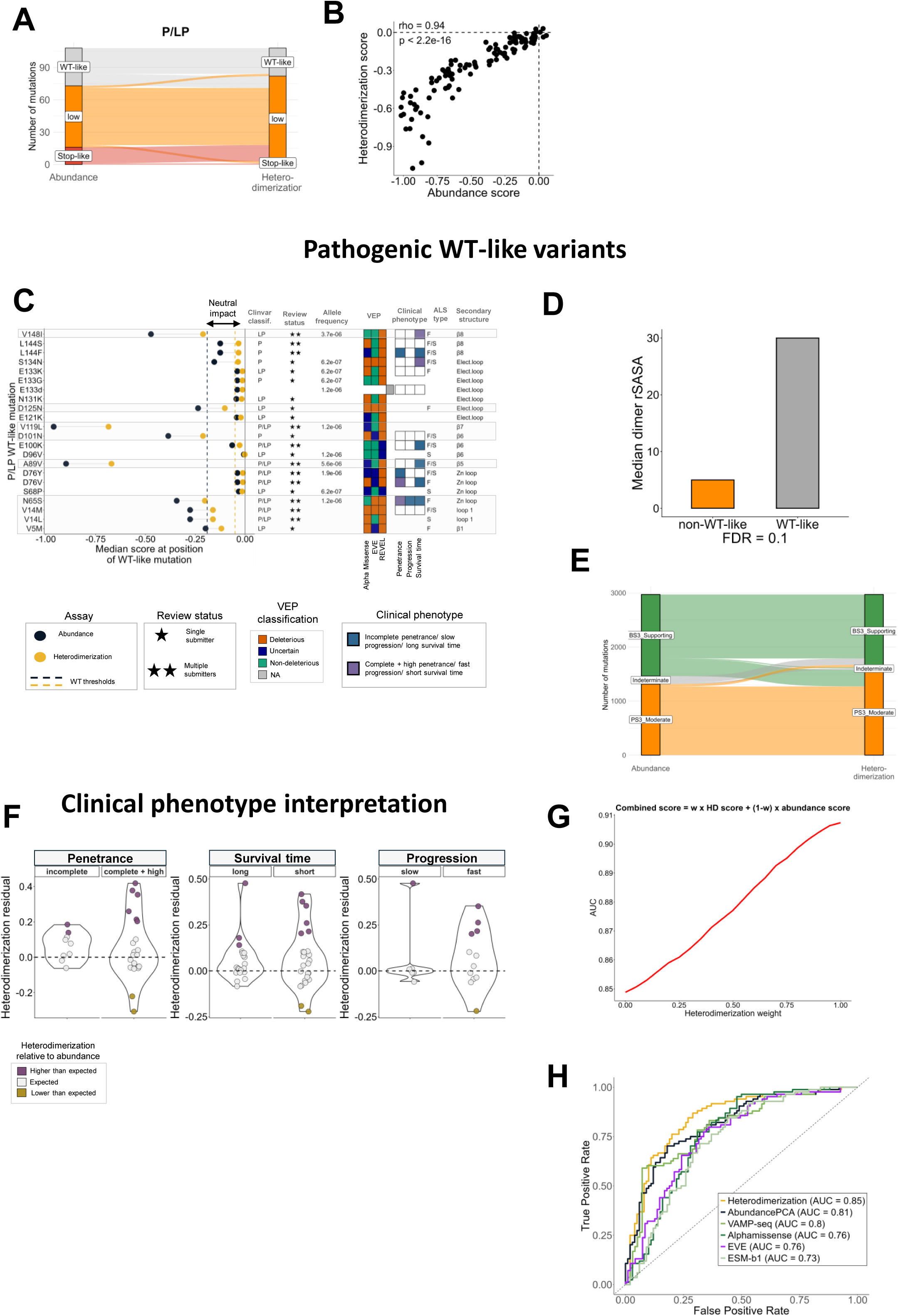
Extended analysis of pathogenic and likely pathogenic SOD1 variants. **(A)** Alluvial plot showing changes in FDR categories between abundance and heterodimerization. **(B)** Scatter plot of the abundance and heterodimerization scores of P/LP variants. Spearman’’s coefficient and p-value are indicated. Dashed lines indicate WT scores. **(C)** Comparison of P/LP WT-like variants features relevant for clinical variant interpretation. From left to right: comparison of each P/LP WT-like (n=22) with the median abundance/heterodimerization score at the position they occur. P/LP variants that occur at positions where the median of scores is lower than the neutral impact range are highlighted in grey; the VEP column indicates the classification of Alphamissense, EVE and REVEL; the review status column indicates the strength of evidence for each WT-like P/LP based on the number of submitters; the clinical phenotype column indicates mild and aggressive phenotypes available for P/LP WT-like variants; the ALS type column indicate whether the variant is involved in sporadic ALS (S), familial ALS (F) or both (F/S); the secondary structure column indicates the region of SOD1 where P/LP variants occur. **(D)** Comparison of the rSASA in the SOD1 dimeric state of variants destabilizing in both abundance and heterodimerization (non-WT-like) and WT-like variants. **(E)** Alluvial plot showing the BS3/PS3 evidence of all SOD1 substitutions of the library in both abundance and heterodimerization. **(F)** Distribution of HD residuals for disease penetrance, survival time and disease progression. Dots are coloured according to HD residual categories. **(G)** Area Under the Curve (AUC) as a function of the weight assigned to abundance and heterodimerization. **(H)** ROC showing the performance of abundance, heterodimerization, VAMP-seq and VEPs at classifying pathogenicity.

**Supplementary figure 10.**
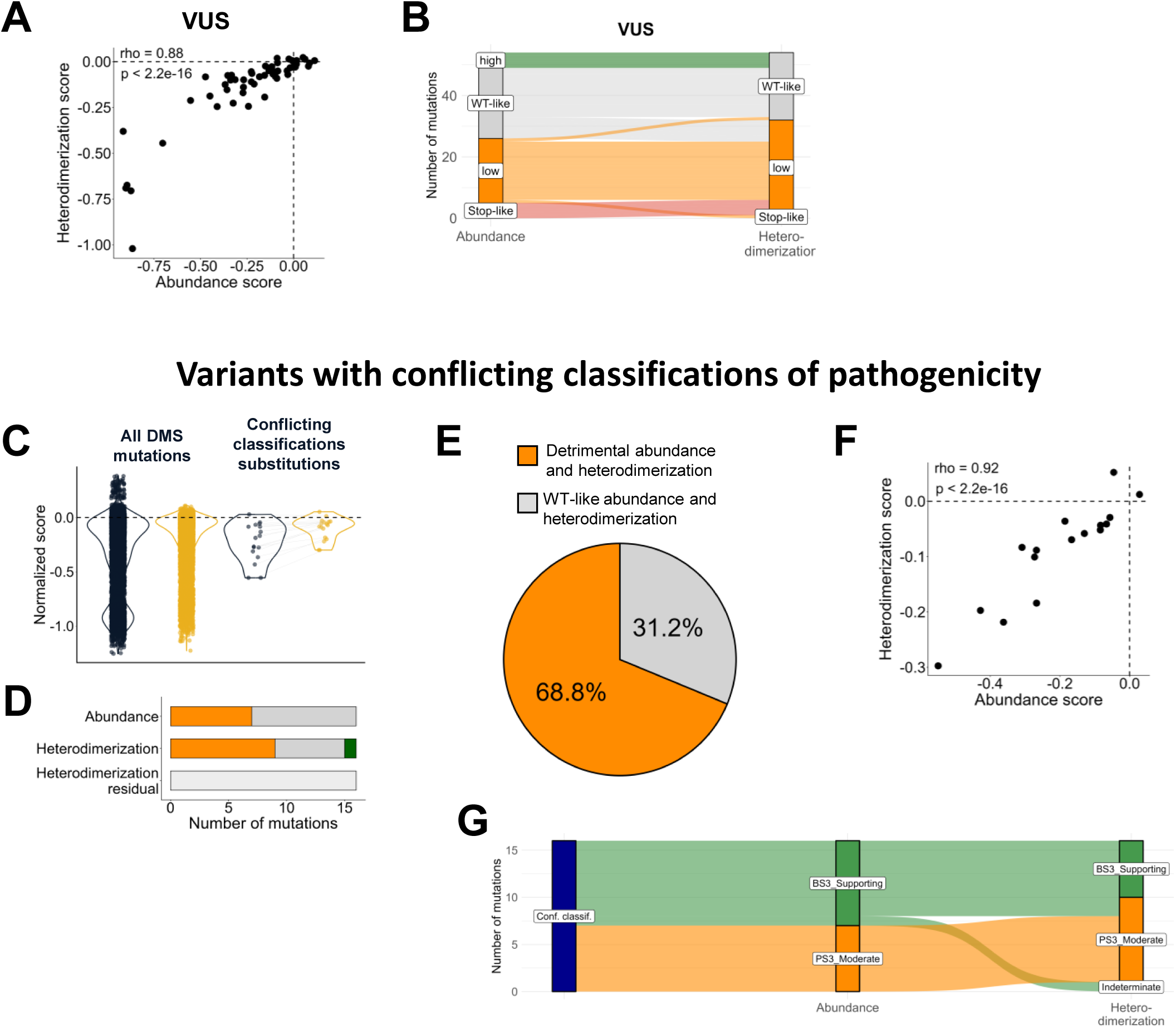
Extended analysis of SOD1 VUS. **(A)** Scatter plot of the abundance and heterodimerization scores of VUS. Spearman’s coefficient and p-value are indicated. Dashed lines indicate WT scores. **(B)** Alluvial plot showing changes in FDR categories between abundance and heterodimerization. **(C)** Violin plot showing the abundance/heterodimerization scores of all the mutations measured in this work (substitutions and indels), and the reported variants with conflicting classifications of pathogenicity. **(D)** Number of conflict. classif. variants with each FDR category based on abundance, heterodimerization, and HD residuals categories. **(E)** Proportion of conflict. classif. variants with low/stop-like classification (orange) and WT-like classification (grey) in both abundance and heterodimerization. **(F)** Scatter plot of the abundance and heterodimerization scores of conflict. classif. variants. Pearson’s coefficient and p-value are indicated. Dashed lines indicate WT scores. **(G)** Alluvial plot showing the BS3/PS3 evidence of conflict. classif. variants in both abundance and heterodimerization.

## Notes

https://github.com/BEBlab/SOD1-ddPCA

### Competing Interest Statement

The authors have declared no competing interest.

### Summary of Updates

Authors list order was updated. There are no other differences.

