## Supplementary material for "The folding, dimerization and allosteric landscapes of the ALS protein SOD1: a comprehensive mutational atlas": Description Supp Files

**Description of Additional Supplementary files**

File Name: Dataset S1

Description: List of oligonucleotides used in this study

File Name: Dataset S2

Description: Description: Processed data required to reproduce the analysis and figures in this paper, with read counts, abundance and heterodimerization scores and associated errors

File Name: Dataset S3

Description: 475 nm (AcGFP) laser transmission value or range used for each FLIM control and SOD1 variant in HEK293TΔSOD1 cells.

File Name: Dataset S4

Description: Individual SOD1 variants measurements with FLIM.

File Name: Dataset S5

Description: Individual SOD1 variants inclusions, survival and intensity GFP measurements.

File Name: Dataset S6

Description: FDR abundance and heterodimerization categories of SOD1 reported variants.

File Name: Dataset S7

Description: Calibration framework of SOD1 variants. It includes the list of pathogenic and proxy benign controls and the LR calculations.

File Name: Dataset S8

Description: SOD1 clinical phenotypes collected from the literature
